# Kinetic gating by an iron–sulfur chain modulates flavin radical-pair spin chemistry in mitochondrial Complex I

**DOI:** 10.64898/2026.10.30.763004

**Authors:** Ji-Yong Sung, Jae-Ho Cheong

## Abstract

Mitochondrial Complex I generates reactive oxygen species at its flavin cofactor, where spin-dependent reaction pathways have been proposed. Models of this chemistry treat the flavin– oxygen radical pair in isolation, although the flavin is wired to a chain of eight iron–sulfur clusters. Here we model the pair with this chain using the human Complex I structure and open-quantum-system models of spin coupling and electron transfer. In the simulated network, spin coherence decays rapidly along the chain, whereas proximal clusters alter the calculated radical-pair singlet yield. Their influence is kinetically gated: a downstream cluster affects the modelled yield when its electron-transfer rate competes with radical-pair recombination, a threshold robust to driving force and to the transfer model. Modelling does set the sign: one-directional spin hops predict suppression and inflate coherence transport, whereas charge-resolved transfer drives the singlet yield towards its spin-randomized limit unless triplet recombination dominates. Detectable spin-dependent effects in this framework require superoxide spin relaxation to be substantially slower than in free solution, a condition not yet established for Complex I.

## Introduction

Mitochondrial Complex I (NADH:ubiquinone oxidoreductase) is the largest and most complex enzyme of the respiratory chain, and its flavin mononucleotide (FMN) cofactor is widely regarded as a principal site of superoxide and hydrogen peroxide production in mitochondria. [1–3] Dysregulated Complex I-derived ROS production has been implicated in ischaemia–reperfusion injury [4] and Parkinson’s disease, [5] as well as in the broader biology of mitochondrial ageing, making the molecular determinants of FMN reactivity a question of substantial physiological and clinical interest.

We recently proposed [6], using a structure-informed radical-pair model of the FMNH•–O2•− pair (the neutral flavin semiquinone and superoxide), that the geometric accessibility of molecular oxygen around the FMN isoalloxazine ring gives rise to a localized maximum of a composite score combining oxygen accessibility with magnetic-field and dephasing responses at near-contact (∼3.3 Å) separation. In extending that analysis we identified three points that we correct here. First, its steric test counted O positions that overlap FMN itself as accessible; once these are excluded, almost no accessible positions remain within 3.3 Å of the isoalloxazine centroid and the composite-score maximum moves to 6–6.5 Å (Fig. 3a and Supplementary Note 2), so the near-contact maximum reported previously does not hold. Second, the earlier code implemented the opposite overall sign of the dipolar coupling term. Third, the reference hyperfine coupling carried forward from that work (aHF = 1.233 mT) describes the anionic FMN•− radical rather than the neutral FMNH• semiquinone, for which the corresponding value is 1.108 mT. Neither of the last two corrections alters any conclusion of the present work (Methods). For brevity, we refer to FMNH• as "FMN" and to O •− as "O " throughout. That analysis, in common with the great majority of radical-pair mechanism studies in spin biology, treated the reacting pair as an isolated two-spin system, with environmental effects folded into a single phenomenological dephasing rate. [7–9] This is a reasonable first approximation, but it leaves an explicit limitation unaddressed: FMN in Complex I is not spin-isolated. [10] It sits at the head of a wire of eight iron–sulfur clusters (N3, N1a, N1b, N4, N5, N6a, N6b, N2) that conducts electrons from FMN to the ubiquinone-binding site over about 95 Å. Six of these are [4Fe–4S] clusters and two are [2Fe–2S] clusters, which can carry unpaired electron spin in appropriate redox states and lie 11–17 Å from their nearest neighbours in the chain — distances at which magnetic dipolar coupling between paramagnetic centres is not obviously negligible. [11]

**Fig. 1.**
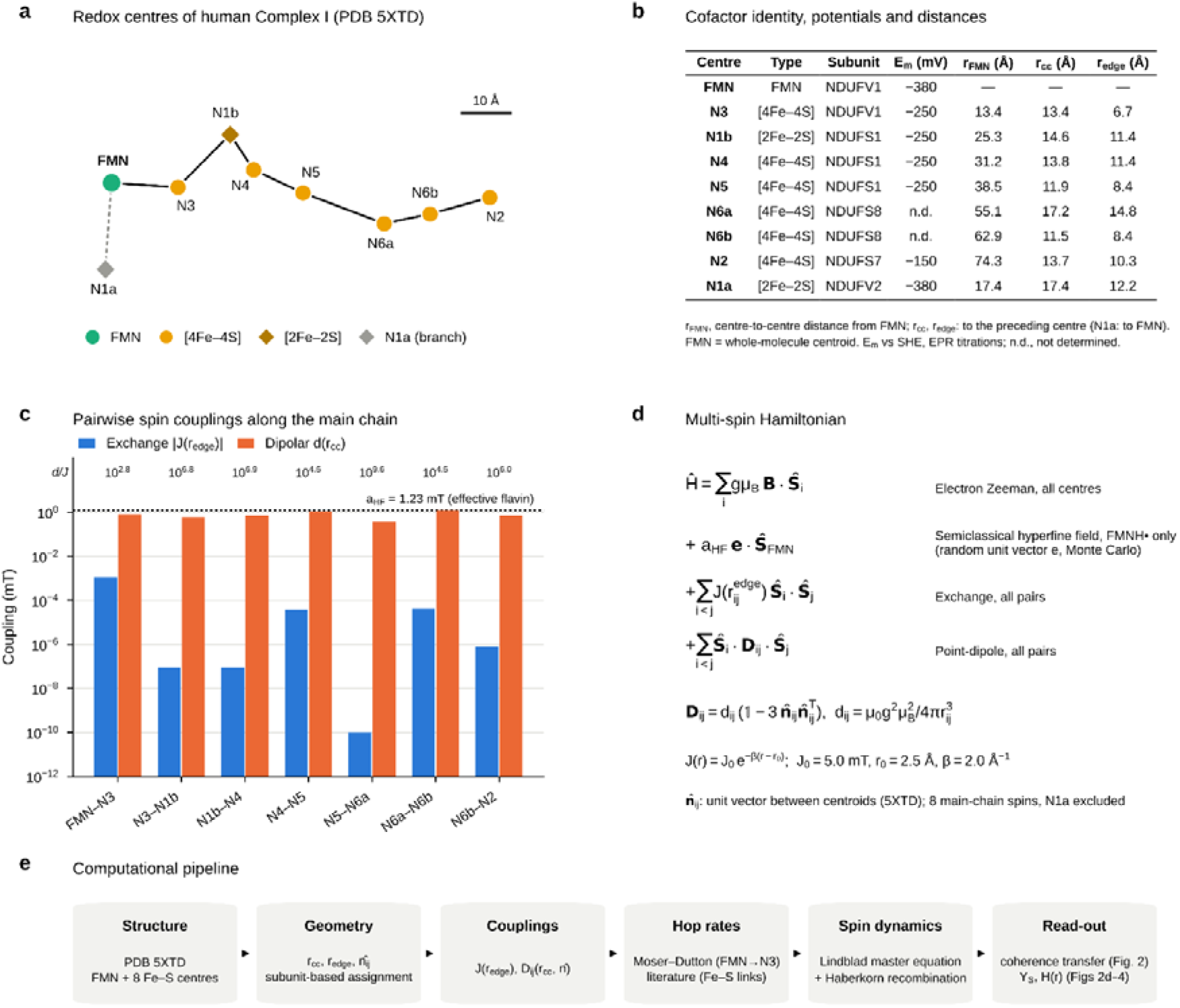
Structure-guided multi-spin network model of the human Complex I electron-transfer chain. a, Redox centres of human Complex I (PDB 5XTD), projected onto the plane of the two largest principal axes of their centroids. The main chain is drawn in electron-transfer order, FMN–N3–N1b–N4–N5–N6a–N6b–N2. FMN is shown in green, [4Fe–4S] clusters as circles and [2Fe–2S] clusters as diamonds. The side-branch cluster N1a (grey, dashed line) is excluded from the main pathway. Apparent distances are shortened by the projection; true distances are given in b. Scale bar, 10 Å. b, Cofactor identity, host subunit, midpoint potential (Em, mV versus SHE, from EPR redox titrations of the mammalian enzyme; n.d., not determined) and distances. rFMN is the centre-to-centre distance from FMN. rcc and redge are the centre-to-centre and edge-to-edge distances to the preceding centre in the chain (for N1a, to FMN). Each centre is located at the unweighted centroid of its cluster iron and sulfur atoms, or of all FMN heavy atoms. Edge-to-edge distance is the minimum heavy-atom distance between two cofactors. c, Exchange coupling (blue) and point-dipole coupling (orange) for each adjacent pair in the main chain, on a logarithmic scale. Exchange was calculated at the edge-to-edge distance as J(r) = J exp[−β(r − r )], with J = 5.0 mT, r = 2.5 Å and β = 2.0 Å ¹. Dipolar coupling was calculated at the centre-to-centre distance as d(r) = μ g²μB²/(4πr³). The ratio d/J is shown above each pair. Dotted line, the effective flavin hyperfine coupling used in the model (1.23 mT). d, Multi-spin Hamiltonian for the eight main-chain spins. It comprises an electron Zeeman term for every centre, a semiclassical hyperfine field on FMNH• only, and exchange and anisotropic point-dipole couplings between all pairs of centres. The dipolar tensor is D = d(1 − 3nn ), where n is the unit vector between centroids in the 5XTD structure. e, Computational pipeline used in this study.

Whether this arrangement constitutes, in any functional sense, a coupled multi-spin network — one whose topology, internode distances and relative orientations could modulate the spin dynamics at the FMN terminus — is an open question that the isolated-pair approximation cannot address. [11, 12] It also sits at the edge of a broader and still-contested problem in spin biology and quantum biology more generally: whether quantum coherence between spatially separated paramagnetic centres in a protein can survive long enough, and propagate far enough, to have any bearing on chemistry occurring elsewhere in the same structure. [13, 14] Most radical-pair mechanism theory to date has been developed and validated for two-spin systems; [15] extending this framework to a multi-cofactor chain requires a qualitatively different theoretical treatment — a many-body open quantum system coupling coherent (exchange, dipolar) and incoherent (electron-hopping, spin relaxation) dynamics simultaneously — and, to our knowledge, has not been attempted for any member of the respiratory chain. A further, long-standing objection concerns superoxide itself: free O • relaxes on a sub-nanosecond timescale because of its unquenched orbital angular momentum, which would erase the spin correlation of any superoxide-containing pair unless the binding site restricts its motion and lifts the orbital degeneracy; recent work has nevertheless shown that tightly bound flavin–superoxide pairs can remain magnetosensitive when recombination is strongly asymmetric. [16] A third body that removes one partner of a radical pair, as proposed for radical scavengers in cryptochrome magnetoreception, can also reshape its spin chemistry [17] [18] [19].

Here we extend our structure-informed spin-dynamics framework from the isolated FMN– O pair to a structure-based eight-site spin network and smaller unified radical-pair models of human Complex I, using atomic coordinates from a cryo-EM structure of the enzyme (PDB 5XTD) together with literature electron-transfer parameters for the iron–sulfur chain. [20, 21] We ask, specifically, how far a spin coherence initialized at one cofactor propagates along the chain before being lost to the fast spin-lattice relaxation characteristic of iron–sulfur clusters, whether the redox state of the chain’s distal terminus (N2, adjacent to the ubiquinone-binding site) has any detectable influence on FMN, [22] and whether coupling FMN to its nearest neighbours in the chain measurably shifts the FMN–O singlet yield and, if so, what sets the size and sign of that shift. Either a negative or a positive answer is informative: robust decoherence within one or two hops would place the isolated-pair approximation on firmer mechanistic footing, whereas measurable network-mediated modulation would motivate treating the Complex I electron-transfer chain as an extended, structurally tunable spin circuit rather than a sequence of independent hops. We find that the chain lies between these alternatives, and that two principal findings emerge across the model comparisons examined. First, the influence of a downstream cofactor is kinetically gated: it switches on, without resonance, once electron transfer to it competes with radical-pair recombination, independently of the driving force and of how transfer is represented. Second, the representation of electron transfer decides the sign of the effect and the apparent reach of spin coherence. A one-directional spin-hop operator, a simple way of adding hopping to a spin Hamiltonian, pumps spin polarization and overstates coherence transport, whereas charge-resolved transfer drives the singlet yield towards its incoherent, spin-randomized limit whenever singlet recombination is at least as fast as triplet recombination.

## Results

### A structure-guided multi-spin network model of the Complex I electron-transfer chain

To extend our previously reported FMN–O radical-pair model beyond the isolated two-spin approximation, we first constructed a structure-guided multi-spin representation of the entire Complex I electron-transfer relay, comprising FMN and its eight associated iron–sulfur clusters (**Fig. 1**). Atomic coordinates for the flavin cofactor and each cluster were extracted directly from a cryo-EM structure of human Complex I (PDB 5XTD), and each redox centre was represented by the centroid of its heavy atoms (**Fig. 1a**). Cluster identities were assigned by tracing the electron-transfer topology from FMN and were independently confirmed by their host subunits (NDUFV1, NDUFS1, NDUFS8, NDUFS7 and NDUFV2). This yielded a nine-node network whose main pathway — FMN–N3–N1b–N4–N5–N6a–N6b–N2 — recapitulates the established electron-transfer sequence of the enzyme, with N1a resolved as a spatially distinct side branch (17.4 Å from FMN) rather than a member of the principal chain (**Fig. 1b**). Centre-to-centre separations along the main pathway ranged from 11.5 to 17.2 Å (mean 13.7 Å) and edge-to-edge separations from 6.7 to 14.8 Å (**Supplementary Table 1**), in close agreement (within 0.6 Å at every step) with independently reported centre-to-centre distances for the homologous Thermus thermophilus enzyme, indicating that chain geometry is essentially conserved across species.

Each node was assigned its cluster type and midpoint potential (**Fig. 1b**): six [4Fe–4S] clusters (N3, N4, N5, N6a, N6b, N2) and two [2Fe–2S] clusters (N1b, N1a), with potentials taken from EPR redox titrations of the mammalian enzyme where available. We then computed, for every adjacent pair in the chain, the isotropic exchange coupling J(r), evaluated at the edge-to-edge separation, and the point-dipole coupling d(r), evaluated at the centre-to-centre separation; all dynamical calculations use the full anisotropic dipolar tensor with the 5XTD inter-centre orientation vector (**Fig. 1c**). Across all seven adjacent pairs, the dipolar coupling exceeded the exchange coupling by roughly three to ten orders of magnitude (d/J = 6 × 10² to 4 × 10 ), and its magnitude (0.36–1.24 mT) was of the same order as the reference FMN hyperfine coupling (aHF = 1.23 mT). This indicates that, to the extent spin coupling propagates along the chain at all, dipolar rather than exchange interactions are expected to dominate its physical origin — a point of direct relevance to the coherence-propagation calculations that follow.

These pairwise couplings were assembled into a single multi-spin Hamiltonian spanning the eight main-chain electron spins (N1a excluded), comprising a Zeeman term for every node, a semiclassical hyperfine term restricted to the FMN node, and exchange and anisotropic dipolar coupling terms between all pairs of centres, parameterized with the real 5XTD distances and orientations (**Fig. 1d**). This Hamiltonian, together with electron-hopping rates obtained from the Moser–Dutton relation and, where available, literature quantum-chemically calculated rates, defines the computational pipeline used throughout this study (**Fig. 1e**): structural extraction, pairwise coupling-constant calculation, hop-rate estimation, many-body Lindblad spin propagation incorporating both coherent (exchange, dipolar) and incoherent (electron-hop, spin-relaxation) channels, and evaluation of the FMN–O singlet yield and its sensitivities.

### A structure-derived, many-body spin model reveals limited but non-zero coherence propagation along the Fe–S chain

Having established the network topology and coupling architecture (**Fig. 1**), we next asked whether spin coherence initialized at FMN propagates measurably along the iron–sulfur chain, and whether perturbations at the chain’s distal terminus (N2, adjacent to the ubiquinone-binding site) can be detected back at FMN. We constructed a multi-spin Lindblad model spanning all eight main-chain nodes (Hilbert-space dimension 2 ), incorporating exchange and anisotropic dipolar coupling between all pairs of centres (**Fig. 1d**) and incoherent electron-hopping channels between adjacent centres, represented by unital flip-flop operators in which each hop is accompanied by its reverse at the same rate, parameterized by literature and Moser–Dutton rates. The three hop rates not available in the literature (N4→N5, N6a→N6b and N6b→N2) were bracketed between 0 and 10 s ¹. Site-resolved relaxation rates reflected the established contrast between the slowly relaxing FMN semiquinone and the much faster-relaxing iron–sulfur clusters (**Methods**). As a precaution against signal generated without injected excitation, which one-directional hop operators produce (**Supplementary Note 1**), all reported trajectories were baseline-subtracted against a control in which every centre starts maximally mixed; as the dynamics are linear, this subtraction was performed exactly by propagating the difference between the two initial states.

Coherence initialized at FMN decayed steeply along the chain (**Fig. 2a, b**). After 5 ns, the coherence remaining at FMN was 4.5 × 10 (from 0.5 at injection), that at N3 was 5.8 × 10 and every more distal centre carried less than 6 × 10 . Values at FMN, N3 and N1b were unaffected by the unknown hop rates, and those at N4–N2 changed by less than 40% between the two bracketing choices. Conversely, coherence injected at the distal terminus, N2, produced a negligible signal at FMN: the maximum within 5 ns decreased monotonically from 2.1 × 10 to 3.0 × 10 as the N2 dephasing rate was increased from 10 to 10¹ s ¹, and changed by about 1% when the unknown hop rates were set to 10 s ¹ (**Fig. 2c**). Together, these results indicate that the Complex I electron-transfer chain does not function as a long-range coherent conduit: the great majority of any propagating spin coherence, and any influence the distal terminus might exert on FMN, is confined to the first one or two nodes of the chain. With the one-directional operators used for the radical-pair calculations below (**Figs 2d and 3**), the same calculations overstated every value: the coherence remaining at FMN after 5 ns was 0.12, and the maximum reaching FMN from N2 was 900–1,000 times larger, because the one-directional operator pumps spin polarization and generates coherence even from a maximally mixed start (**Supplementary Fig. 5** and **Supplementary Note 1**).

**Fig. 2.**
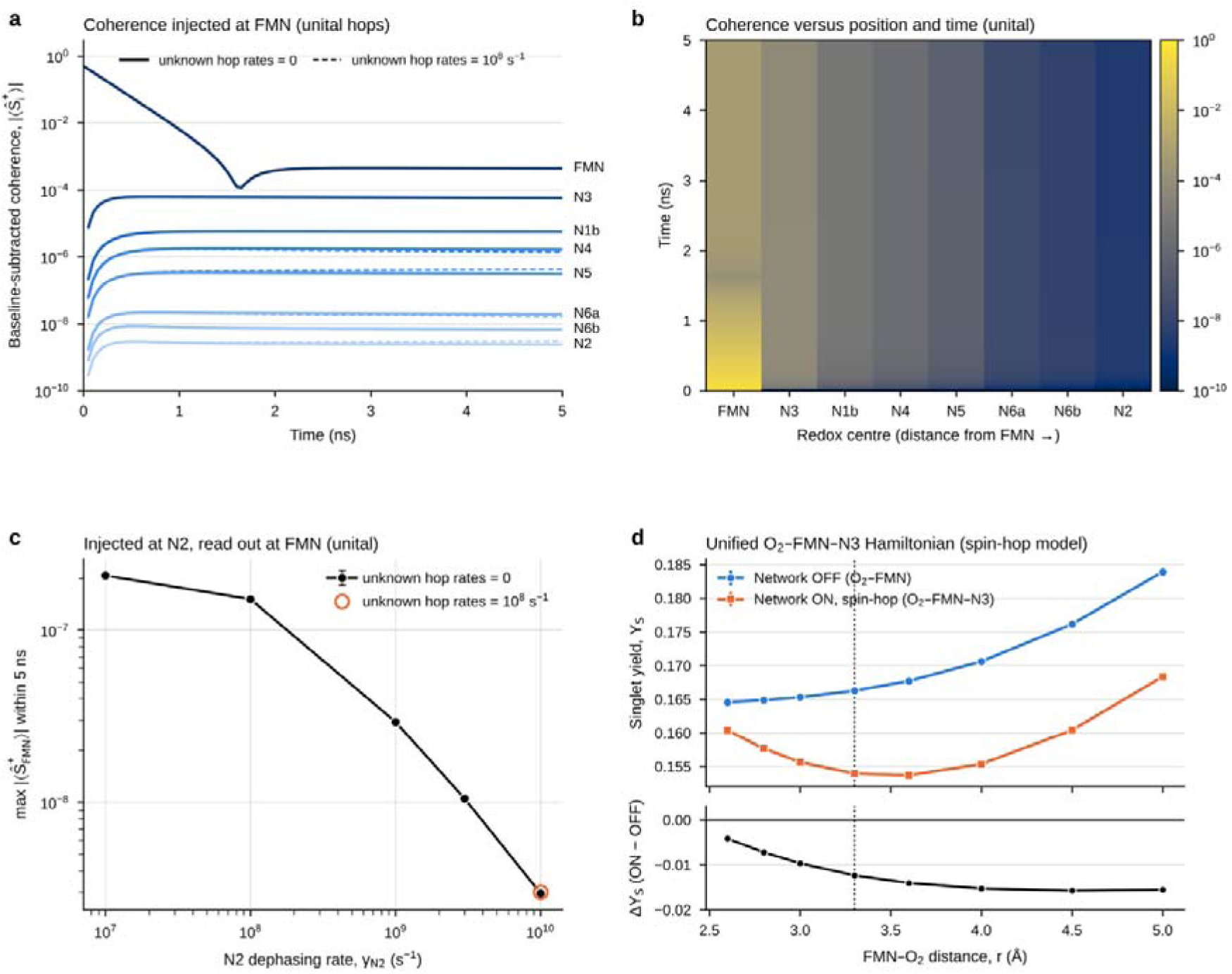
Limited coherence propagation along the Fe–S chain and (in the spin-hop model) suppression of the FMN–O singlet yield by the nearest network node at slow radical-pair recombination. a, Node-resolved transverse spin coherence, |⟨S ⟩|, after coherence is injected at FMN. The calculation used an eight-site Lindblad model (FMN, N3, N1b, N4, N5, N6a, N6b and N2; Hilbert-space dimension 2 ). The model includes an electron Zeeman term at every centre (0.05 mT) and a semiclassical hyperfine field on FMNH• (1.233 mT). It also includes all-pairs exchange and anisotropic point-dipole couplings from the PDB 5XTD geometry (Fig. 1), pure dephasing (3 × 10 s ¹ for FMN and 10¹ s ¹ for each Fe–S cluster) and unital electron-hop channels between adjacent centres (each flip-flop accompanied by its reverse at the same rate). Traces are baseline-subtracted: each is the excited trajectory minus a control in which every centre starts maximally mixed. Values are averages over 12 random hyperfine orientations. Solid lines, the three hop rates not available in the literature (N4→N5, N6a→N6b and N6b→N2) set to zero; dashed lines, these rates set to 10 s ¹. The dashed lines are shown only where they differ from the solid lines (N4–N2). After 5 ns, coherence was 4.5 × 10 at FMN, 5.8 × 10 at N3 and below 6 × 10 at every further centre. The dip in the FMN trace near 1.6 ns marks a change in phase of the ensemble-averaged coherence. Results with the one-directional operators of Figs 2d and 3 are compared in Supplementary Fig. 5. b, Heat map of the data in a (solid lines) as a function of centre position and time. c, Terminal transfer function. Coherence was injected at N2, and the plot shows the maximum baseline-subtracted coherence reaching FMN within 5 ns as a function of the dephasing rate assumed for N2. The transfer decreases monotonically from 2.1 × 10 to 3.0 × 10 as the N2 dephasing rate increases from 10 to 10¹ s ¹. Error bars, s.e.m. Open circle, unknown hop rates set to 10 s ¹, which changed the transfer by 1.4%. d, Singlet yield (YS) of the FMN–O radical pair as a function of FMN–O separation. YS was computed from a unified O –FMN–N3 Hamiltonian (Hilbert-space dimension 2³) with Haberkorn recombination (kS = kT = 10 s ¹). Network OFF is the isolated O –FMN pair. Network ON adds FMN–N3 exchange and dipolar coupling from the 5XTD geometry, dephasing at N3 and an FMN→N3 hop channel (8.5 × 10 s ¹, Moser–Dutton relation). Yields were evaluated exactly at infinite time and averaged over 400 random hyperfine orientations, with the same orientations used for both conditions. The bottom panel shows the paired difference, ΔYS = YS(ON) − YS(OFF). ΔYS is negative at every separation (−0.0042 to −0.0157; signal-to-noise ratio 39–182). Dashed vertical line, the separation of the composite-score maximum in our earlier analysis (3.3 Å). Error bars, s.e.m.; where not visible, they are smaller than the symbols.

**Fig. 3.**
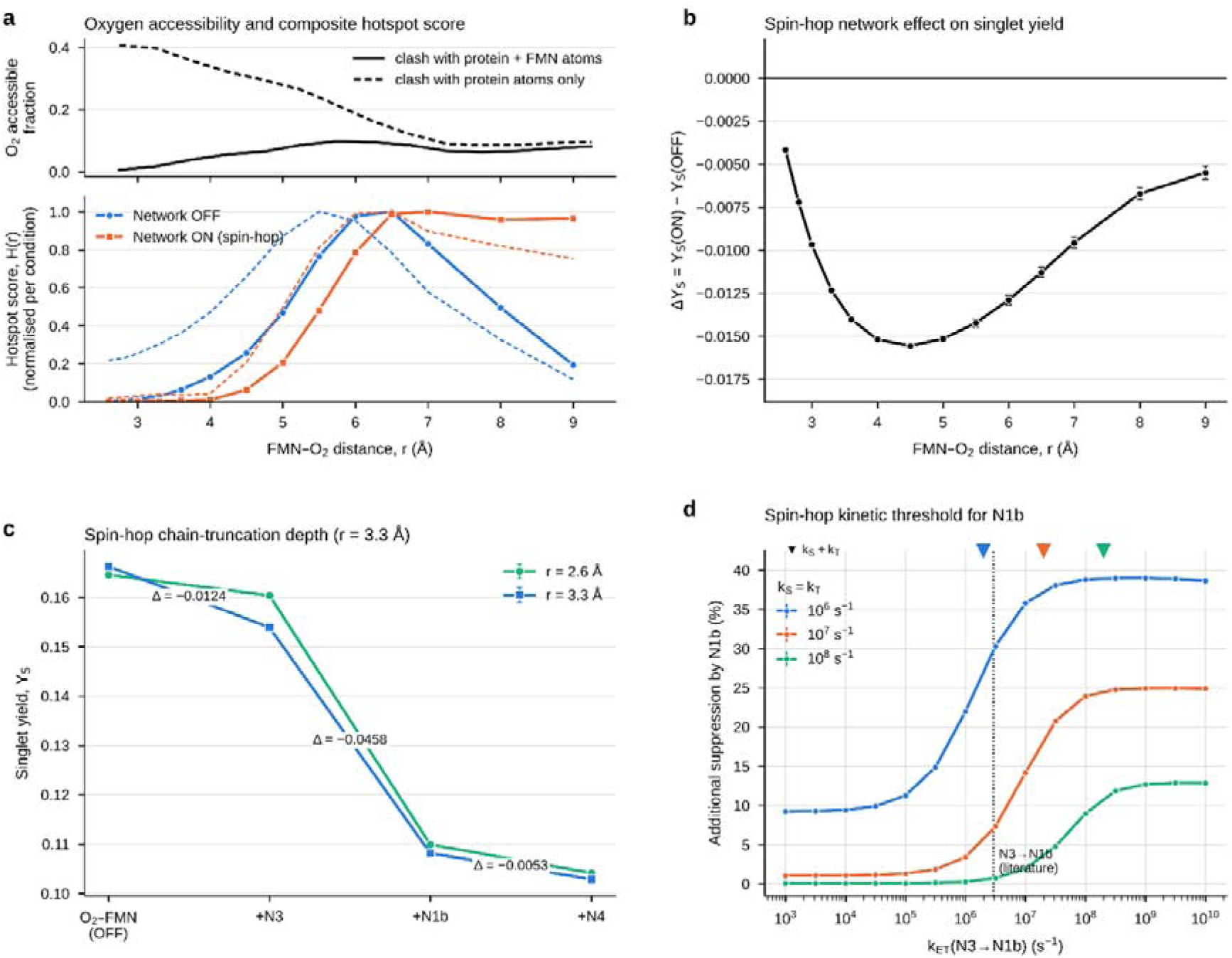
In the phenomenological spin-hop model, network coupling suppresses the singlet yield at slow recombination, and the N1b contribution follows a kinetic threshold. a, Top, fraction of candidate O positions that are sterically accessible, as a function of distance from the FMN isoalloxazine centroid. Candidate positions were 5 × 10 points sampled uniformly within a 15 Å sphere in PDB 5XTD and binned in 0.5 Å shells. Solid line, a position counts as accessible only if no protein heavy atom lies within 3.0 Å and no FMN heavy atom lies within 2.8 Å. Dashed line, protein atoms only are tested, with FMN atoms omitted. Bottom, composite hotspot score, H(r) = A(r) × ½[MB(r) + Cγ(r)], for network OFF (blue) and network ON in the spin-hop model (orange). A(r) is the normalised accessibility, MB(r) the magnetic-field sensitivity of the singlet yield and Cγ(r) its dephasing sensitivity. Solid lines use the accessibility that includes FMN atoms; dashed lines use protein atoms only. Each curve is normalised to its own maximum. The ratio of the combined spin sensitivities is shown in Supplementary Fig. 4a. b, Paired difference in singlet yield, ΔYS = YS(ON) − YS(OFF), across the sampled distance range (unified O –FMN–N3 spin-hop model). ΔYS is negative at every separation (signal-to-noise ratio 15–415). Error bars, s.e.m. c, Singlet yield as the spin-hop Hamiltonian is extended stepwise from the isolated O –FMN pair to include N3, N1b and N4, at FMN–O separations of 2.6 Å (green) and 3.3 Å (blue). Numbers give the change in YS at each step for r = 3.3 Å. d, Additional suppression of the singlet yield contributed by N1b in the spin-hop model, as a function of the N3→N1b electron-hop rate, expressed as [YS(+N3) − YS(+N3+N1b)] / YS(+N3), for kS = kT = 10 s ¹ (blue), 10 s ¹ (orange) and 10 s ¹ (green), at r = 3.3 Å. Triangles on the top axis mark kS + kT; dotted line, the literature N3→N1b rate (2.9 × 10 s ¹). The contribution rises sigmoidally and saturates, with no resonance maximum, and the midpoint moves with the reaction rate (0.25–0.7 × (kS + kT)); for r = 6.5 Å see Supplementary Fig. 2b, and for the charge-resolved model Fig. 4b. Yields were evaluated exactly at infinite time with kS = kT = 10 s ¹ unless stated and averaged over random hyperfine orientations (2,000 in a and b, 200 in c, 150 in d), with the same orientations used in every compared condition. Magnetic-field sensitivity was obtained by finite difference at 0.05 ± 0.02 mT, and dephasing sensitivity between 1.5 × 10 and 6 × 10 s ¹. Error bars, s.e.m.; where not visible, they are smaller than the symbols.

### In a phenomenological spin-hop model, a unified O –FMN–N3 Hamiltonian predicts that the nearest network node suppresses the FMN–O singlet yield at slow recombination

The limited spatial reach of coherence propagation motivated a tractable, non-reduced test of whether the nearest network node influences the FMN–O radical pair directly. We constructed a single, unified Hamiltonian spanning O , FMN and N3 (**Fig. 2d**), in which the preceding paper’s radical-pair formalism — Haberkorn recombination kinetics and exponential exchange coupling, supplemented here with an axial O –FMN point-dipole term — was combined in one quantum master equation with the real-geometry FMN–N3 exchange and anisotropic dipolar coupling and the corresponding electron-hopping channel (represented, unlike the unital hops of the eight-site model, by a one-directional spin flip-flop jump operator, √k σ σ ; we refer to this as the spin-hop model and test it against charge-resolved electron transfer in Fig. 4), rather than being connected post hoc through an effective parameter. Singlet yields were evaluated exactly at infinite time and compared between this network-ON condition and the isolated O –FMN pair (network OFF) using the same 400 random hyperfine orientations for both. At these baseline recombination rates (kS = kT = 10 s ¹), network coupling lowered the singlet yield at every FMN–O separation between 2.6 and 5.0 Å (ΔYS = −0.0042 to −0.0157; signal-to-noise ratio 39–182); at 3.3 Å, the separation of the composite-score maximum in our earlier analysis, YS fell from 0.166 to 0.154. We retain 3.3 Å as a reference separation for continuity with that analysis, although it lies within the sterically excluded region identified below (**Fig. 3a**); the distance dependence of each effect is reported explicitly (**Figs 3b** and **4e**). We report this and the following spin-hop results because a one-directional jump operator is a simple way of adding electron hopping to spin models, and because they reveal the kinetic gating analysed next (**Fig. 3**). Fig. 4 shows that this gating survives a charge-resolved treatment, whereas the direction of the effect at slow recombination does not.

**Fig. 4.**
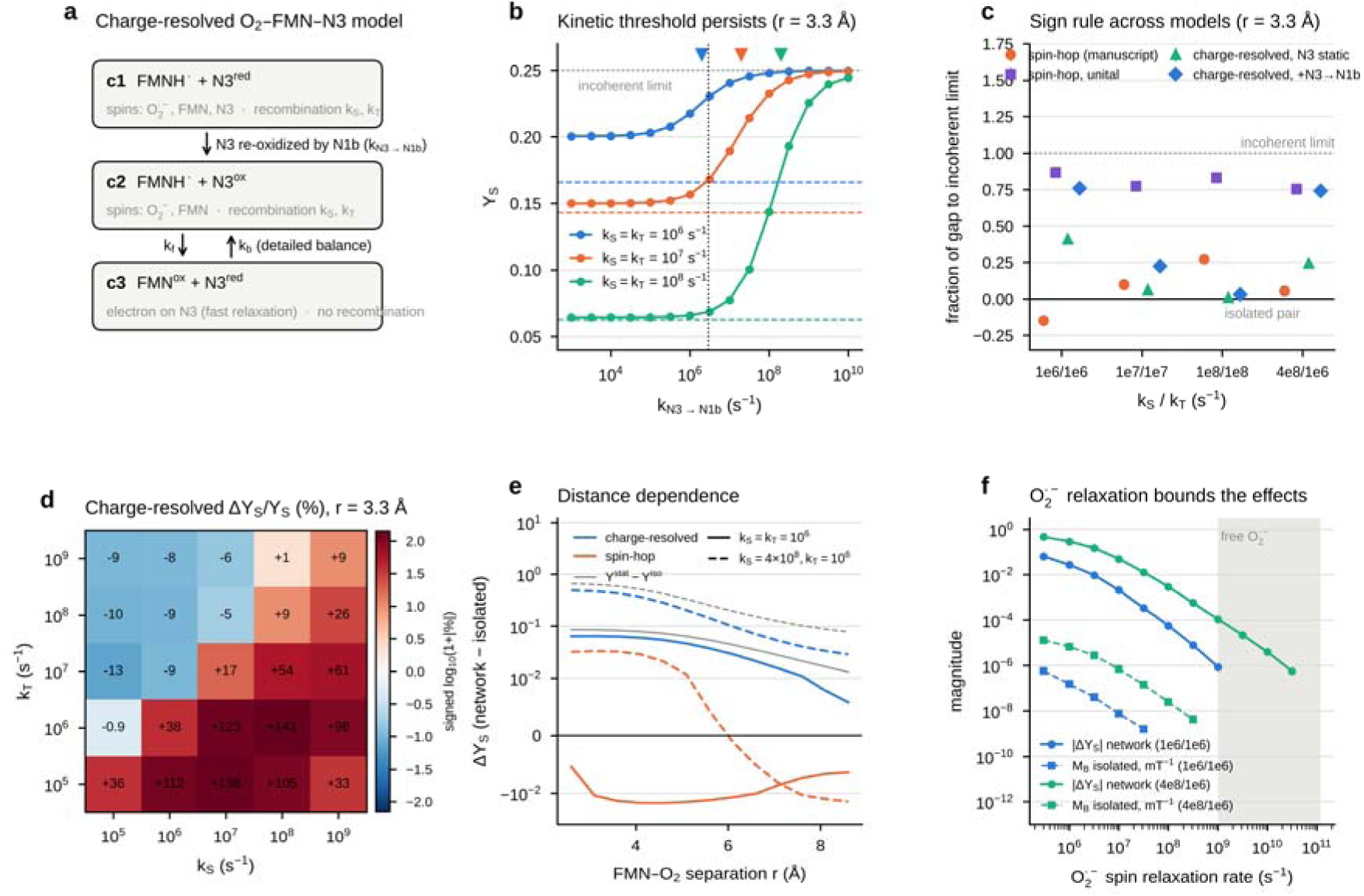
Charge-resolved electron transfer preserves the kinetic threshold and, for kS ≥ kT, the iron–sulfur chain drives the FMN–O radical pair towards its incoherent limit. a, Charge-resolved O –FMN–N3 model. The system occupies one of three redox configurations: c1, FMN semiquinone with N3 reduced; c2, FMN semiquinone with N3 oxidized (diamagnetic); c3, oxidized FMN with the transferred electron on N3, which cannot recombine with O • . N3 is re-oxidized by N1b (c1→c2); FMN N3 transfer is spin-conserving and obeys detailed balance at 310 K (kf = 8.5 × 10 s ¹, kb = 2.9 × 10 s ¹). Electrons on N3 relax isotropically (10¹ s ¹). Calculations start from the reduced chain (c1). b, Singlet yield as a function of the N3→N1b re-oxidation rate at r = 3.3 Å for kS = kT = 10 (blue), 10 (orange) and 10 s ¹ (green). Dashed lines, isolated pair; dotted horizontal line, incoherent limit kS/(kS + 3kT) = 1/4; dotted vertical line, literature N3→N1b rate (2.9 × 10 s ¹); triangles, kS + kT. The rise is sigmoidal with no resonance maximum, and its midpoint (0.6–1.0 × (kS + kT)) moves with the reaction rate. 150 paired hyperfine orientations per point. c, Fraction of the gap between the isolated-pair yield and the incoherent limit closed by network coupling, (YS − YS^iso^)/(YS^stat^ − YS^iso^), at r = 3.3 Å for four recombination regimes, in the one-directional spin-hop model used in Figs 2d and 3 (orange), the same model made unital by adding the reverse flip-flop at equal rate (purple), and the charge-resolved model with N3 held reduced (green) or re-oxidized by N1b at the literature rate (blue). Negative values indicate motion away from the incoherent limit. 400 paired orientations; error bars, s.e.m. d, Percentage change in YS produced by the charge-resolved network at r = 3.3 Å across a grid of recombination rates (numbers in cells; colour, signed logarithm). The sign agrees with that of YS^stat^ − YS^iso^ at all 15 points with kS ≥ kT; at kT > kS, where both yields are below 0.035, the network lowers YS by 1–13% at nine of ten points, moving it away from the incoherent limit. 150 paired orientations per point. e, Network-induced change in YS as a function of FMN–O separation for the charge-resolved (blue) and spin-hop (orange) models, at kS = kT = 10 s ¹ (solid) and kS = 4 × 10 , kT = 10 s ¹ (dashed); grey, YS^stat^ − YS^iso^. Symmetric-logarithmic axis (linear within ±0.01). f, Effect of O • spin relaxation (isotropic, rate kO2) on the magnitude of the charge-resolved network effect (solid) and on the low-field magnetic sensitivity of the isolated pair, MB (dashed), at r = 3.3 Å for kS = kT = 10 s ¹ (blue) and kS = 4 × 10 , kT = 10 s ¹ (green). Points at the left edge, no O • relaxation beyond the baseline dephasing. Shaded, sub-nanosecond relaxation expected for free superoxide. All yields were evaluated exactly at infinite time with matched hyperfine orientations across compared conditions.

### Oxygen accessibility places the spin-sensitivity hotspot at 6–6.5 Å

We next asked how coupling FMN to its neighbouring iron–sulfur clusters alters the spin sensitivity of the FMN–O pair across the binding pocket. We computed the magnetic-field and dephasing sensitivities of the singlet yield for network OFF and ON, and combined them with oxygen accessibility into the composite hotspot score H(r) (**Fig. 3a**). Accessibility was recalculated from the 5XTD structure by Monte Carlo sampling with a steric test that excludes positions overlapping protein or FMN atoms. Under this test, positions within 3.3 Å of the isoalloxazine centroid were almost entirely inaccessible (≤ 2%), and accessibility peaked near 6 Å. In both conditions, the magnetic-field sensitivity was very small at near-contact separations, where strong O –FMN exchange and dipolar coupling dominate the spin dynamics, and rose more than 100-fold by 6 Å. As a result, H(r) peaked at 6.0–6.5 Å for network OFF (values within 2% of each other) and reached a broad plateau at 6.5–9 Å for network ON (within 5% of its maximum), and neither condition showed a near-contact maximum; this held even when FMN atoms were omitted from the steric test (peaks at 5.5 and 6.5 Å). In the spin-hop model, network coupling also increased the combined spin sensitivity, by up to 35-fold and entirely through its magnetic-field term (**Supplementary Fig. 4a**); the charge-resolved model does not reproduce this increase (below).

### The network suppresses singlet yield throughout the sampled FMN pocket at slow radical-pair recombination in the spin-hop model

To determine whether this enhanced sensitivity corresponds to a change in reactive outcome, we computed the singlet yield, YS, directly from the same unified Hamiltonian across the full sampled distance range (r = 2.6–9.0 Å; **Fig. 3b**). Network coupling lowered YS at every distance tested, with no sign reversal: ΔYS was −0.0042 at 2.6 Å, reached its largest magnitude (−0.0156) at 4.5 Å and fell to −0.0055 at 9.0 Å (2,000 paired hyperfine orientations; signal-to-noise ratio 15–415). The network therefore raises the spin sensitivity of the pocket (**Supplementary Fig. 4a**) while uniformly lowering the singlet yield: the effect of coupling on local responsiveness and its effect on reaction outcome are dissociable quantities.

### In the spin-hop model, suppression grows with the first additional network node

Because the unified Hamiltonian above included only the nearest cluster (N3), we asked how the suppressive effect scales as further nodes are added to the chain. Extending the Hamiltonian stepwise to include N1b and then N4 (**Fig. 3c**) showed that the suppression of YS does not saturate at N3. At r = 3.3 Å, adding N3 lowered YS by 0.012, adding N1b lowered it by a further 0.046, and adding N4 by only 0.005; the same pattern held at 2.6 Å (0.004, 0.050 and 0.006). The N1b step was thus 4–12-fold larger than the N3 step and about ninefold larger than the N4 step. The small N4 step is consistent with the kinetic picture developed below: the N1b→N4 rate (6.4 × 10 s ¹) is about 30-fold slower than the radical-pair reaction rate (kS + kT = 2 × 10 s ¹). Extension to N5 was not attempted because no literature rate constant is available for the N4→N5 step. Accordingly, the network-ON quantities in **Fig. 3a,b**, which use the N3-only Hamiltonian, should be interpreted as lower bounds on the magnitude of the suppression; their direction is re-examined with a charge-resolved model below (**Fig. 4**).

### The N1b contribution reflects a kinetic threshold set by the radical-pair reaction rate

The disproportionate N1b step coincides with a near-equality between the N3→N1b hop rate (2.9 × 10 s ¹) and the radical-pair reaction rate (kS + kT = 2 × 10 s ¹). To test whether this reflects a resonance, we varied the N3→N1b rate from 10³ to 10¹ s ¹ at three reaction rates (kS = kT = 10 , 10 and 10 s ¹; **Fig. 3d**). At this separation, in every case, the additional suppression contributed by N1b rose sigmoidally with the hop rate and saturated above about 10 s ¹, with no maximum at rate matching. The midpoint of the rise moved with the reaction rate (1.3 × 10 , 8.2 × 10 and 5.0 × 10 s ¹, respectively, or 0.25–0.7 × (kS + kT)). The N1b contribution therefore reflects a kinetic threshold rather than a resonance: a downstream cofactor alters the radical-pair outcome once electron transfer to it competes with recombination, and faster transfer produces the largest effect.

The large N1b step at the baseline reaction rate arises because the literature N3→N1b rate lies on the rising edge of this threshold, which makes the N1b contribution strongly dependent on the assumed reaction rate. At the literature hop rate, the additional suppression by N1b was about 30% of YS at kS = 10 s ¹, 7% at 10 s ¹ and below 1% at 10 s ¹ (**Fig. 3d**).

In the spin-hop model the direction of the nearest-neighbour effect also depends on the recombination kinetics and on the FMN–O separation (**Supplementary Figs 1 and 2** and **Supplementary Note 5**); we revisit this dependence with the charge-resolved model below.

The spin-hop yields were insensitive to the FMN→N3 rate over 10 –10¹¹ s ¹ (**Supplementary Fig. 4b**), and at the baseline rates the pair had essentially fully reacted within 6 µs (**Supplementary Fig. 4c**), which is shorter than the slowest hop included in the extended Hamiltonians (N1b→N4, τ ≈ 16 µs). All effects reported here are therefore established on the timescale of a single radical-pair reaction; multi-turnover dynamics were not probed.

### Charge-resolved electron transfer preserves the kinetic threshold but reverses the sign at slow recombination

The one-directional spin-hop operator of Figs 2d and 3 moves spin between a fixed set of paramagnetic centres without creating or removing any, and, because it acts in one direction only, it is non-unital: it continuously pumps spin polarization onto FMN. To test whether our conclusions depend on this representation, we built a charge-resolved model in which the O –FMN–N3 system occupies one of three redox configurations (**Fig. 4a**): c1, the FMN semiquinone with N3 reduced (three electron spins); c2, the semiquinone with N3 oxidized and diamagnetic (two spins); and c3, oxidized FMN with the transferred electron on N3, which cannot recombine with O • (two spins). The transferred electron carries its spin, FMN→N3 transfer is reversible with detailed balance at 310 K (kf = 8.5 × 10 s ¹, kb = 2.9 × 10 s ¹), N3 is re-oxidized by N1b (c1→c2) at the N3→N1b rate, and electrons on N3 relax isotropically at 10¹ s ¹ (**Methods**).

The redox state of the chain determines whether a radical pair can persist at all. When N3 was initially oxidized and the transferred electron could move on to N1b, the semiquinone electron left FMN within about 0.1 ns and the pair was effectively quenched (YS = 0.003 at kS = kT = 10 s ¹; **Supplementary Fig. 3b**), consistent with the proposal that the electron remaining on the flavin radical after superoxide formation passes to the iron–sulfur clusters [1]. This quenching depends on slow recombination: at kS = 4 × 10 , kT = 10 s ¹ the same calculation gave YS = 0.53, above the isolated-pair value, and with slower onward transfer (6.4 × 10 s ¹) YS at kS = kT = 10 s ¹ ranged from 0.18 to 0.002 as the FMN→N3 driving force was varied from −0.05 to −0.20 eV (**Supplementary Note 6**). Under the slow-recombination conditions of this model, an initially reduced N3 permits substantially greater radical-pair persistence than an oxidized N3, the condition under which Complex I produces the most superoxide [1], and we used the reduced chain (c1) as the initial configuration below.

Starting from the reduced chain, coupling to N3 raised rather than lowered the singlet yield. At 3.3 Å and kS = kT = 10 s ¹, YS increased from 0.166 for the isolated pair to 0.200 with N3 held reduced and to 0.230 when N3 could be re-oxidized by N1b at the literature rate (400 paired orientations; signal-to-noise ratio > 1,000; **Fig. 4c**), compared with 0.153 in the spin-hop model. Varying the N3→N1b rate recovered the kinetic threshold of Fig. 3d (**Fig. 4b**): YS rose sigmoidally from its static-N3 value towards 1/4, with no maximum, and the midpoint moved with the reaction rate (2.0 × 10 , 1.6 × 10 and 1.3 × 10 s ¹ at kS = kT = 10 , 10 and 10 s ¹, or 0.6–1.0 × (kS + kT); 0.7–2.2 × (kS + kT) at 6.5 Å). At the literature N3→N1b rate the gate was 59%, 18% and 2% open at these three reaction rates. The kinetic threshold is therefore not an artefact of the spin-hop representation, but the direction of the effect at slow recombination is. None of these reduced-chain yields changed by more than 10 when the FMN→N3 driving force was varied from −0.05 to −0.20 eV (kf = 5.2 × 10 –2.6 × 10¹ s ¹; kb = 8.1 × 10 –1.5 × 10 s ¹), a range that brackets the uncertainty in the one-electron flavin potential: once N3 is re-oxidized, FMN N3 shuttling is much faster than recombination for every choice (Supplementary Note 6). At the sterically accessible separation of 6.5 Å, the yields were 0.214 for the isolated pair, 0.226 with N3 held reduced and 0.236 with N3 re-oxidizable at kS = kT = 10 s ¹, and 0.833, 0.856 and 0.908 at kS = 4 × 10 , kT = 10 s ¹.

### With charge-resolved transfer, the chain drives pairs with kS ≥ kT towards their incoherent limit

Making the spin-hop operator unital, by adding the reverse flip-flop at the same rate, was enough to reverse its sign: at 3.3 Å and slow recombination YS became 0.239 instead of 0.153 (**Fig. 4c**). The suppression in Figs 2d and 3 thus arises from the spin polarization pumped onto FMN by the one-directional operator rather than from electron transfer as such. (**Supplementary Note 4**). At these rates, in both charge-resolved configurations and the unital spin-hop control examined (the unital spin-hop model and the charge-resolved model with N3 held reduced or re-oxidized), coupling to the chain moved YS from its isolated-pair value, YS^iso^, towards the incoherent limit YS^stat^ = kS/(kS + 3kT), the yield of a triplet-born pair whose spin correlation is fully randomized before it reacts (**Fig. 4c**).

Across a 5 × 5 grid of recombination rates (10 –10 s ¹ each), the charge-resolved effect had the sign of YS^stat^ − YS^iso^ at all 15 points with kS ≥ kT, increasing YS by 9–141% (**Fig. 4d**). Where triplet recombination dominates (kT > kS), both yields are small (YS < 0.035) and at nine of ten points the chain lowered YS by 1–13% even though the isolated pair lay below the incoherent limit, moving the yield away from that limit. The tendency towards the incoherent limit is therefore not a general rule; it holds for pairs with kS ≥ kT, a regime proposed for a flavin–superoxide pair. [16] It also accounts for the distance dependence (**Fig. 4e**). In the charge-resolved model the chain raised YS at every FMN–O separation from 2.6 to 8.6 Å, both at kS = kT = 10 s ¹ and at kS = 4 × 10 , kT = 10 s ¹, closing 37–76% of the gap to the incoherent limit; the sign change with distance seen in the spin-hop model at fast recombination (**Supplementary Fig. 2a**) did not occur.

The chain therefore acts as a kinetically gated spin-randomizing sink: a fast-relaxing reduced N3 and, more strongly, reversible charge shuttling after re-oxidation of N3 erode the coherent singlet–triplet interconversion of the pair. The gated part of the effect was insensitive to the poorly known Fe–S relaxation rate: with N3 re-oxidizable, YS at 3.3 Å varied only between 0.226 and 0.230 for rates from 10 to 10¹² s ¹, whereas the contribution of a static reduced N3 peaked near 10¹ s ¹ and vanished in the fast-relaxation (motional-narrowing) limit (**Supplementary Fig. 3a**).

### Superoxide spin relaxation bounds the network and magnetic-field effects

Free superoxide relaxes on a sub-nanosecond timescale, so we asked how much O • spin relaxation the network effect tolerates (**Fig. 4f**). Isotropic O • relaxation drove both the isolated and the network-coupled pair towards the incoherent limit and erased the network effect: at 3.3 Å, the charge-resolved change in YS fell from 0.064 without additional relaxation to 0.002 at 10 s ¹ and below 10 at 10 s ¹ for kS = kT = 10 s ¹, and from 0.46 to 0.003 at 10 s ¹ and 10 at 10 s ¹ for kS = 4 × 10 , kT = 10 s ¹. The low-field magnetic sensitivity of the isolated pair decayed in parallel. Under the adopted relaxation model and recombination rates, appreciable network and magnetic-field effects require bound superoxide spin relaxation below approximately 10 –10 s ¹, at least an order of magnitude more slowly than free O • ; whether the Complex I pocket immobilizes superoxide to this extent, as proposed for cryptochrome, [16] is unknown. The charge-resolved network also did not reproduce the enhanced magnetic sensitivity of the spin-hop model (Supplementary Fig. 4a). At 6.5 Å it lowered MB, by 2.3-fold at kS = kT = 10 s ¹ and by 25-fold at kS = 4 × 10 , kT = 10 s ¹, whereas the spin-hop network raised it by more than an order of magnitude (**Supplementary Fig. 3c**), as expected if the chain randomizes rather than polarizes the FMN spin.

## Discussion

Building on our earlier model of the FMN semiquinone–superoxide radical pair in Complex I treated in isolation, we asked whether this description survives contact with the eight-cluster iron–sulfur relay to which FMN is structurally and electronically coupled. The answer is neither a simple "yes" nor a simple "no." Spin coherence initialized at FMN decays by several orders of magnitude within one or two hops along the chain (**Fig. 2**), and a perturbation at the chain’s distal terminus has a correspondingly negligible effect on FMN (**Fig. 2c**) — properties consistent with the fast spin-lattice relaxation known to characterize iron–sulfur clusters at physiological temperature. In this narrow sense, the isolated-pair approximation used in our preceding analysis captures the dominant physics: Complex I is not a long-range coherent quantum wire connecting the ubiquinone site to FMN.

Yet coupling FMN to even its nearest neighbour is not inconsequential, and how it matters depends on how electron transfer is represented. In the phenomenological spin-hop model the chain suppresses the singlet yield at slow recombination (**Figs 2d, 3b**), with the largest contribution from the N3→N1b link (**Fig. 3c**). A charge-resolved treatment, in which electron transfer creates and removes paramagnetic centres and obeys detailed balance, keeps the central kinetic result: the influence of a downstream cofactor rises sigmoidally once electron transfer to it competes with recombination, saturates when transfer is much faster and shows no resonance (**Figs 3d, 4b**). It reverses the sign at slow recombination, however, and shows why: the chain is a spin-randomizing sink that pushes the singlet yield towards its incoherent limit, kS/(kS + 3kT), whereas the one-directional spin-hop operator adds a spurious spin-polarizing drive (**Fig. 4c**). For kS ≥ kT, the regime expected for a flavin–superoxide pair, the chain moved YS towards the incoherent limit and, because coherent evolution left the isolated pair below that limit in every such condition we examined, raised it (**Fig. 4d,e**). This is a tendency rather than a general rule: where triplet recombination dominates, the chain moved the yield away from the limit (**Fig. 4d**).

The comparison also carries a methodological lesson for spin-dynamics models of other electron-transfer chains, such as the tryptophan triad of cryptochrome or photosynthetic reaction centres. A one-directional spin-hop operator is a convenient way to add hopping to a spin Hamiltonian, but it is non-unital. Here it reversed the sign of the network effect at slow recombination (**Fig. 4c**), inflated the low-field magnetic sensitivity by more than an order of magnitude (**Supplementary Fig. 3c**) and, in the eight-site model, overstated the coherence reaching FMN from the distal terminus by about three orders of magnitude (**Supplementary Fig. 5**). The kinetic threshold survived every representation, whereas the sign and reach of the effect did not. Electron transfer between radicals should therefore be represented as the movement of charge, with paramagnetic centres created and removed and detailed balance obeyed, or at least by unital operators, before conclusions are drawn about the direction or range of network effects.

The charge-resolved model connects the Complex I chain to the radical-scavenging mechanism proposed for cryptochrome magnetoreception, in which a third radical that reacts with one partner of a radical pair reshapes, and can amplify, its magnetic sensitivity [17][18][19]. In Complex I the scavenger is built into the protein, and its effective rate is set by the redox state and electron-transfer kinetics of the chain: at slow recombination an oxidized N3 quenches the pair within nanoseconds, whereas a reduced N3 acts through its fast-relaxing spin and, after re-oxidation by N1b, through reversible charge shuttling. This suggests two experimentally testable, conditional predictions. First, slowing an electron-transfer step below the radical-pair reaction rate, for example by mutating residues on its tunnelling pathway, should largely remove that cofactor’s contribution, whereas accelerating it should not increase the contribution beyond saturation. Second, because the radical pair requires a reduced chain, spin-dependent changes in the modelled flavin radical-pair yield are most likely under reducing conditions (high NADH/NAD ), and, for kS ≥ kT, their direction should follow the incoherent limit once kS and kT are known. Both predictions presuppose that bound superoxide relaxes more slowly than about 10 –10 s ¹ (**Fig. 4f**).

A second observation concerns where in the pocket the radical pair is most sensitive. In the isolated pair, magnetic-field sensitivity is negligible at near-contact separations, where strong O –FMN exchange and dipolar coupling dominate, and the composite sensitivity score peaks at 5.5–6.5 Å (**Fig. 3a**). This supersedes the near-contact maximum reported in our earlier analysis. The difference arises mainly from the steric test. Positions that overlap FMN itself, which were counted as accessible in our earlier analysis, are now excluded, and almost no accessible positions remain within 3.3 Å of the isoalloxazine centroid (**Fig. 3a**). When the earlier calculation is repeated with its own definitions, excluding these positions alone moves the composite-score maximum from near contact to beyond 4 Å, whereas none of the other differences between the two models (hyperfine representation, O –FMN dipolar term, sensitivity definitions or normalisation) does so (**Supplementary Note 2 and Supplementary Table 2**). In the spin-hop model, network coupling increases the combined spin sensitivity at every resolved separation, by up to 35-fold through its magnetic-field component (**Supplementary Fig. 4a**), while lowering the singlet yield everywhere at the baseline rates (**Fig. 3b**). Local responsiveness to perturbation and the direction of the chemical outcome are therefore dissociable properties of a coupled spin network: a region of high sensitivity is not necessarily one in which coupling favours a particular reactive channel. The charge-resolved model, however, does not reproduce the sensitivity enhancement and instead lowers the low-field magnetic sensitivity (**Supplementary Fig. 3c**), so this enhancement, like the suppression, should be attributed to the polarizing spin-hop operator. The location of the accessibility-weighted hotspot, which is set by the isolated pair and the steric test, is not affected.

These findings should be read against several explicit constraints. First, our tractable unified Hamiltonians include at most three network nodes (**Fig. 3c**); the full chain was not treated. Second, most calculations were performed at r = 3.3 Å for continuity with our earlier analysis, although this separation is sterically disfavoured (**Fig. 3a**); the charge-resolved network effect keeps its sign at every separation from 2.6 to 8.6 Å (**Fig. 4e**), whereas the spin-hop effect changes sign with distance at fast recombination (**Supplementary Fig. 2a**). Third, all reported effects are established on the timescale of a single radical-pair reaction (the pair has essentially fully reacted within 6 µs at the baseline rates), which is shorter than the slowest hop in the extended Hamiltonians (N1b→N4, τ ≈ 16 µs) and the chain’s slowest electron-transfer step (N5→N6a, τ ≈ 137 µs); multi-turnover dynamics were not probed. Fourth, rate constants for three Fe–S links (N4→N5, N6a→N6b and N6b→N2) are not available; bracketing them between 0 and 10 s ¹ changed coherence at the distal centres by less than 40% but did not alter any conclusion. Fifth, the charge-resolved model includes only FMN, N3 and the re-oxidation of N3 by N1b; it treats the clusters as g ≈ 2 point spins at their centroids with isotropic relaxation, ignoring the delocalization of cluster spin density over the iron sites, and neglects O • –N3 coupling. Moreover, re-oxidation of N3 by N1b at the literature rate presupposes an oxidized N1b, whereas the baseline slowly recombining radical pair persists preferentially with reduced N3 in the present model; when the whole chain is reduced, re-oxidation would instead be limited by slower downstream steps. The NADH-reduced enzyme retains a partially oxidized cluster ensemble [27], and the calculation with N3 held reduced (**Fig. 4c**) brackets the opposite extreme, but the relevant redox populations remain to be determined. Anisotropic g tensors, explicit vibronic electron-transfer theory and a longer charge-resolved chain would refine the magnitudes; the kinetic threshold persisted over the tested parameter ranges (FMN→N3 driving force, Fe–S relaxation rate and exchange convention), whereas the direction of the effect depends on the recombination regime (**Fig. 4d**). Sixth, O • spin relaxation in the Complex I pocket is unknown; network and magnetic-field effects are bounded by it (**Fig. 4f**) and vanish if bound superoxide relaxes as fast as free superoxide. Even without such relaxation, the low-field magnetic sensitivity of the isolated pair is small in absolute terms (|ΔYS/ΔB| ≤ 10 ³ mT ¹ at 6.5 Å; **Fig. 4f**). Finally, we used numerically exact propagation, which is adequate for Hilbert spaces of up to 2 dimensions but not for the full chain, for which tensor-network methods would be appropriate.

Several extensions follow from these constraints. Obtaining rate constants for the three missing links, extending the unified Hamiltonian beyond N4 and treating the N1a branch cluster explicitly would test whether the network effect saturates. The response of the network-coupled pair to local structural fluctuations, which amplified the isolated-pair response in our earlier work, also remains to be examined. Our treatment of relaxation uses phenomenological pure dephasing rather than the Nakajima–Zwanzig relaxation superoperator recently applied to a closely related flavin radical pair in cryptochrome, [16] which also captures the quantum Zeno regime of strongly asymmetric recombination. Given the likely asymmetry between kS and kT for an O -containing pair, the Complex I site may sit closer to that regime than assumed here, and adopting this framework is a natural next step.

More broadly, these results suggest that biological electron-transfer chains harbouring paramagnetic cofactors should not be assumed, a priori, to behave either as fully coherent quantum networks or as sequences of independent, mutually invisible radical pairs. The Complex I relay sits between these extremes: locally coupled and globally decoherent, with its influence on radical-pair chemistry set by which nearby cofactors receive electrons faster than the pair recombines, and, for kS ≥ kT, with a direction set by the incoherent limit of the isolated pair. Because this kinetic threshold reflects competition between electron transfer and recombination in these models, analogous gating may occur in other multi-centre redox enzymes, and our framework is positioned to test this in other systems. Testing this will require representing electron transfer as the movement of charge, because the one-directional spin-hop shortcut reversed the predicted effect under the conditions tested.

## Methods

### Structure preparation and cofactor identification

Atomic coordinates for FMN and the eight associated iron–sulfur clusters of human Complex I were extracted from the cryo-EM structure deposited as PDB entry 5XTD. [21] HETATM records corresponding to FMN, SF4 ([4Fe–4S]) and FES ([2Fe–2S]) ligands were parsed using Biopython, [23] yielding nine redox centres: one FMN (chain A, residue 502), one [4Fe–4S] cluster in the same subunit (chain A, residue 501), five further [4Fe–4S] clusters (chains B, C and M) and two [2Fe–2S] clusters (chains M and O). Each cluster’s geometric centroid was computed as the unweighted mean position of its constituent heavy atoms. FMN was represented by the centroid of all its heavy atoms.

Redox-centre identity was assigned as follows. N1a was identified as the [2Fe–2S] cluster in NDUFV2 (chain O), the only cofactor in that subunit, and treated as a side branch (17.4 Å from FMN). The main chain was then traced from FMN by greedy nearest-neighbour traversal, appending at each step the unassigned centre nearest to the current chain terminus, which yielded the sequence N3, N1b, N4, N5, N6a, N6b and N2. [10] The assignment was confirmed by host subunit: FMN and N3 in NDUFV1 (chain A), N1b, N4 and N5 in NDUFS1 (chain M), N6a and N6b in NDUFS8 (chain B) and N2 in NDUFS7 (chain C). Adjacent-pair centre-to-centre distances agreed to within 0.6 Å with those reported for the homologous Thermus thermophilus enzyme. [24]

Two categories of inter-cofactor distance were computed from the resulting coordinates. Centre-to-centre distances, used for the point-dipole coupling calculation, were taken as the Euclidean distance between cluster centroids. Edge-to-edge distances, used for electron-hop rate estimation, were taken as the minimum pairwise distance between any heavy atom of one cluster and any heavy atom of the adjacent cluster, computed directly from the full HETATM coordinate sets (not approximated from centroid geometry). Unit vectors describing the orientation of each adjacent pair were computed from the centroid-to-centroid displacement, for use in the anisotropic point-dipole Hamiltonian term (below). The O accessibility analysis (below) measures distance from the centroid of the FMN isoalloxazine ring, where the semiquinone spin density resides. [11, 25–27]

### Coupling-constant calculation

#### Exchange coupling

The isotropic exchange coupling between adjacent redox centres was modelled with the exponential distance-decay form established in the preceding two-spin analysis, J(r) = J exp[−β(r − r )], with J = 5.0 mT, r = 2.5 Å and β = 2.0 Å ¹, evaluated at the true edge-to-edge separation for each pair. [28] This functional form and its parameters were reused without refitting; because it was originally calibrated over the 2.5–9 Å range relevant to the FMN–O pair, its extrapolation to the larger (6.7–14.8 Å) edge-to-edge separations spanning the Fe–S chain returns exchange couplings several orders of magnitude below the point-dipole term at every link (**Fig. 1c**) and is used only to confirm that exchange is negligible along the chain. [28]

#### Dipolar coupling

The perpendicular point-dipole coupling was calculated as d(r) = μ g²μB²/(4πr³), expressed in mT, with g = 2.0023 and r the centre-to-centre separation. All multi-spin Hamiltonians (**Figs 2–4**) use the full anisotropic tensor, D = d ( − 3n n ), so that the interaction is Ŝ ·D ·Ŝ = d [Ŝ ·Ŝ − 3(Ŝ ·n )(Ŝ ·n )], with n the 5XTD unit vector connecting the two centres (cryo-EM structure). Tensors were computed from this closed-form expression and verified against RadicalPy. [29] Our earlier code implemented the opposite overall sign of the dipolar term; repeating the unified-Hamiltonian calculations with both signs changed ΔYS by at most 0.0008 and did not alter any conclusion. All distances, unit vectors and coupling constants are listed in **Supplementary Table 1**.

#### Redox potentials and driving forces

The midpoint potentials shown in **Fig. 1b** are EPR redox-titration values for the mammalian enzyme: N3, N1b, N4 and N5, −250 mV; [30] N2, −150 mV; N1a, −380 mV; [20] FMN (two-electron couple), −380 mV. N6a and N6b have not been resolved by titration. [30] The driving force for the FMN→N3 step was computed for the one-electron FMN/FMNH• couple (≈ −340 mV, as in our earlier work) and N3 (−250 mV) as ΔG = −(E°′acceptor − E°′donor)/1000 = −0.09 eV. [31] Because the one-electron potentials of the Complex I flavin are uncertain, the charge-resolved calculations were repeated with ΔG from −0.05 to −0.20 eV (**Supplementary Note 6**). Other steps used literature rate constants (below), so no further driving forces were required. [32]

### Electron-hop rate estimation

Forward electron-transfer rates were taken from quantum-chemical (ZINDO tunnelling-current) calculations for the water-mediated pathway where rate constants are available (N3→N1b, N1b→N4 and N5→N6a). [33] Rate constants for the remaining three Fe–S links (N4→N5, N6a→N6b and N6b→N2) are not available; in the eight-site model they were bracketed between 0 and 10 s ¹, and they do not enter the unified Hamiltonians, which extend only to N4. [33] The FMN→N3 rate was estimated with the Moser–Dutton relation [21], log kET = 15 − 0.6Redge − 3.1(ΔG + λ)²/λ, with λ = 0.5 eV and Redge the 5XTD edge-to-edge distance (6.71 Å). Electron-hop channels were implemented as Lindblad jump operators √kET σ σ between adjacent sites. In the eight-site model each operator was paired with its reverse, √kET σ σ , at the same rate (unital hops); the unified radical-pair models use the one-directional operator alone, and the two are compared in **Supplementary Fig. 5** and **Fig. 4c**. [34]

### Electron-hop rate values

The rate constants used were: FMN→N3, 8.5 × 10 s ¹ (Moser–Dutton); N3→N1b, 2.9 × 10 s ¹; N1b→N4, 6.4 × 10 s ¹; and N5→N6a, 7.3 × 10³ s ¹ (τ ≈ 137 µs). All rate constants and their sources are listed in **Supplementary Table 1**.

### Eight-site coherence-propagation model

The eight main-chain centres (FMN, N3, N1b, N4, N5, N6a, N6b and N2) were each represented as an electron spin-½ (Hilbert-space dimension 2 = 256). [35] The Hamiltonian comprised an electron Zeeman term for every centre (B = 0.05 mT), a semiclassical hyperfine field on FMN (aHF = 1.233 mT) whose orientation was sampled randomly for each Monte Carlo trajectory (repeating the key calculations with 1.108 mT, the value for the neutral flavin dataset, changed no conclusion; [36] **Supplementary Note 3 and Supplementary Table 3**), and exchange and anisotropic dipolar couplings between all pairs of centres, computed from the 5XTD geometry as described above. Dissipation comprised (i) pure dephasing, implemented as √(γ /2) σ z, so that transverse coherence decays at rate γ , with γFMN = 3 × 10 s ¹ and γFe–S = 10¹ s ¹ for every cluster (an order-of-magnitude estimate based on the loss of resolvable Fe–S EPR signals above about 20–25 K, implying sub-nanosecond relaxation at physiological temperature), and (ii) the electron-hop channels described above. [35, 37]

Coherence-propagation trajectories (**Fig. 2a, b**) [38] were obtained by preparing the excited site in |+x and every other site in the maximally mixed state /2, which is appropriate because electron Zeeman splittings at these fields are many orders of magnitude smaller than kBT at 310 K. [39] The control trajectory starts with every site in /2; with unital hops this control is stationary ( ρ mixed ≈ 10 s ¹), whereas with one-directional hops it develops a nonzero coherence at FMN even without excitation (**Supplementary Note 1** and **Supplementary Fig. 5**); the one-directional calculation was repeated with the same 12 orientations for comparison. Because the Lindblad dynamics are linear, the baseline-subtracted signal was obtained exactly by propagating the traceless difference of the two initial states, Ŝx,k ( /2)^ 7, for excitation at site k. Propagation used the sparse Liouvillian with SciPy’s expm_multiply over 0–5 ns (101 time points) and was averaged over 12 random hyperfine orientations. [39] Terminal transfer functions (**Fig. 2c**) were computed by exciting N2 and recording the maximum transverse coherence, | Ŝ |, at FMN within 5 ns for N2 dephasing rates of 10 –10¹ s ¹. [40]

### Unified O –FMN–N(chain) Hamiltonian

To connect the network dynamics to FMN–O radical-pair chemistry within a single quantum master equation, the isolated radical-pair Hamiltonian of our earlier work was combined with the network Hamiltonian in one tensor-product Hilbert space (2 ²- dimensional for N Fe–S centres). [39] The Hamiltonian comprised an electron Zeeman term for all sites; the FMN hyperfine term; exponential exchange and axial point-dipole coupling between O and FMN, swept over r = 2.6–9.0 Å; and, for each included network link, exchange, anisotropic dipolar coupling and an electron-hop channel parameterized from the 5XTD geometry, with dephasing at each included Fe–S cluster. [36] Network OFF is the same code path with all network terms removed. Removing the FMN–N3 exchange term changed YS by less than 0.005% at every r, confirming that exchange is negligible. Pure dephasing was implemented as √(kSTD/2) σ z on O and FMN (kSTD = 3 × 10 s ¹) and as √γFe–S σ z (transverse decay rate 2γFe–S) on each included Fe–S centre (γFe–S = 10¹ s ¹). [37] This transverse rate is twice that of the eight-site model; halving it changed the spin-hop ΔYS from −0.0123 to −0.0142 at 3.3 Å and from −0.0104 to −0.0197 at 6.5 Å, without changing its sign.

### Haberkorn recombination was implemented as a non-Hermitian term, Ĥeff = Ĥ − i(kS/2)P S

− i(kT/2)P T, with P S and P T the singlet and triplet projectors of the O –FMN pair. [41–43] The Liouvillian was = −i(spre[Ĥeff] − spost[Ĥeff†]) + Σ [ĉ ], combining this non-trace-preserving term with the dephasing and electron-hop dissipators. The reference rates kS = kT = 10 s ¹ were carried forward from our earlier work; a diffusion-influenced estimate for a flavin semiquinone–superoxide pair is about 4 × 10 s ¹. The dependence on the recombination rates was examined with an exact kS–kT grid at r = 3.3 Å (10 –10 s ¹ for each rate, 400 paired orientations per point; **Supplementary Fig. 1**) and with the N3→N1b rate sweep in **Fig. 3d**. [44] The distance dependence at fast recombination and the N3→N1b sweep at r = 6.5 Å (**Supplementary Fig. 2**) used 1,000 and 400 paired orientations, respectively. [34, 45] Time integration on grids coarser than the radical-pair lifetime (for example, 5 ns steps at kS = 4 × 10 s ¹) underestimated YS with network coupling by about 16% and reversed the apparent sign of the network effect; the exact expression used here avoids this error. [46]

Singlet yields were evaluated exactly at infinite time as YS = kS Tr[P S(− ) ¹ρ (0)], [44] and finite-window yields (**Supplementary Fig. 4c**) [46] as YS(T) = kS Tr[P S ¹(e^( T) − 1)ρ (0)]. The initial state was the maximally mixed triplet state of the O –FMN pair, P T/3, as in our earlier work, with every network spin in /2. At kS = 10 s ¹, this exact evaluation agrees with time integration of the master equation (QuTiP mesolve with normalize_output = False, 6 µs window) to within 0.05% for the same hyperfine orientation.

Yields were averaged over random hyperfine orientations (400 in **Fig. 2d**; 2,000 in **Fig. 3a,b and Supplementary Fig. 4a**; 200 in **Fig. 3c**; 150–400 in **Fig. 3d and Supplementary Fig. 4b,c**). [47]

Network ON used only the nearest node (O –FMN–N3, 2³ = 8-dimensional) except in the chain-truncation analysis (**Fig. 3c**) and the N3→N1b rate sweep (**Fig. 3d**), in which the Hamiltonian was extended to N1b and N4 (2 - and 2 -dimensional Hilbert spaces, respectively). [33] Extension to N5 was not attempted because the N4→N5 rate constant is unavailable. [33]

### Charge-resolved O –FMN–N3 model

The charge-resolved model (Fig. 4a) is defined on the direct sum of three redox configurations: c1 (FMNH•, N3 reduced; spins O • , FMN and N3; dimension 8), c2 (FMNH•, N3 oxidized; spins O • and FMN; dimension 4) and c3 (FMN oxidized, transferred electron on N3; dimension 4), giving a 16-dimensional Hilbert space. In c1 and c2 the Hamiltonian of the O –FMN pair has the same terms as in the unified model (Zeeman, semiclassical FMN hyperfine field, exponential exchange and axial point-dipole coupling); in c1 it also contains the Zeeman term of N3 and the anisotropic FMN–N3 point-dipole coupling from the 5XTD geometry, and in c3 it contains only Zeeman terms. Exchange enters as −2J Ŝ_O ·Ŝ_FMN rather than the +J Ŝ_O ·Ŝ_FMN of the unified model; using the unified-model convention changed YS by at most 0.011 and changed no sign (**Supplementary Note 6**). Haberkorn recombination of the O –FMN pair acts in c1 and c2 only. Electron transfer is represented by jump operators between configurations that conserve the spin of the transferred electron: FMN→N3 (c2→c3, √kf times the identity on the two spins) and N3→FMN (c3→c2, √kb), with kf = 8.5 × 10 s ¹ (Moser–Dutton) and kb = kf exp(ΔG/kBT) = 2.9 × 10 s ¹ for ΔG = −0.09 eV and T = 310 K; and re-oxidation of N3 by N1b (c1→c2) at the N3→N1b rate, implemented as two operators √k m|N3 (m = ↑, ↓) that trace out the departing electron spin. Onward transfer from c3 (requiring prior re-oxidation of N1b, 6.4 × 10 s ¹) was set to zero in the reduced-chain calculations and varied in Supplementary Fig. 3b. Pure dephasing of O • and FMN is as in the unified model; electrons on N3 (in c1 and c3) relax isotropically through the three operators √(γ/4) σ x,y,z, so that T1 ¹ = T2 ¹ = γ = 10¹ s ¹ unless stated. The initial state was (P T/3) /2 in c1 (reduced chain) or P T/3 in c2 (oxidized N3). Yields were computed exactly as YS = kSTr[P S(− ) ¹ρ (0)], with a regularization of 10 ³ s ¹ added to the diagonal of − to remove the null space of unpopulated coherences (effect < 10 on YS). With kf = 0 and the system started in c2, the model reproduces the isolated-pair yields of the unified code to within 10 when the same exchange convention is used.(**Supplementary Note 4**).

### Unital spin-hop control, incoherent limit and superoxide relaxation

The unital spin-hop control adds, to each one-directional jump operator √k σ iσ j of the unified model, its reverse √k σ jσ i at the same rate. The incoherent limit YS^stat^ = kS/(kS + 3kT) is the singlet yield of a triplet-born pair whose four spin states are equally populated at all times. Spin relaxation of O • was added to the isolated, spin-hop and charge-resolved models as isotropic random-field relaxation, √(kO2/4) σ x,y,z on the O spin, with kO2 = T1 ¹ = T2 ¹ varied from 10 to 10¹¹ s ¹; the magnetic-field sensitivity was evaluated as MB = | YS(0.07 mT) − YS(0.03 mT) |/(0.04 mT). The Fe–S relaxation rate was varied from 10 to 10¹² s ¹ in the same way. Recombination-rate grids, distance sweeps, relaxation sweeps and rate sweeps used 150 paired hyperfine orientations per point, and the model comparison (**Fig. 4c**) used 400.

### Composite hotspot score and structural accessibility (Figure 3a)

Following the form of our earlier composite score, the composite hotspot score was H(r) = Ã(r) × ½[MB (r) + Cγ (r)], where Ã(r) is the normalised oxygen accessibility and MB and Cγ are the magnetic-field and dephasing sensitivities of YS normalised to their network-OFF maxima. The sensitivities were the absolute values of orientation-averaged finite differences, MB = | YS(B + ΔB) − YS(B − ΔB) |/(2ΔB) at B = 0.05 mT and ΔB = 0.02 mT, and Cγ = | YS(2k) − YS(k/2) |/(1.5k) at k = 3 × 10 s ¹, where · denotes the average over hyperfine orientations and every evaluation for a given orientation used the same hyperfine draw. Unlike our earlier analysis, which compared singlet yields at about 0.05 and 50 mT, MB is here a local derivative at geomagnetic field strength. In **Fig. 3a** each H(r) curve is normalised to its own maximum; **Supplementary Fig. 4a** compares network ON and OFF on the common network-OFF normalisation. Accessibility was computed from 5 × 10 points sampled uniformly within a 15 Å sphere centred on the FMN isoalloxazine centroid. A point was counted as accessible if no protein or non-FMN heteroatom heavy atom lay within 3.0 Å and no FMN heavy atom lay within 2.8 Å; as a control, accessibility was also computed with protein atoms only (our earlier analysis used an element-specific van der Waals test against protein atoms). The accessible fraction was binned in 0.5 Å shells.

### Statistical analysis

For all network ON/OFF and parameter-sweep comparisons, Monte Carlo samples were drawn using matched random-number-generator seeds across the compared conditions, so that each condition’s YS estimate used the identical sequence of classical hyperfine-vector orientations; this paired design substantially increases statistical power relative to comparing independently sampled means, because most of the Monte Carlo variance is common to both conditions and cancels in the per-sample difference. Reported uncertainties are standard errors of the mean (s.e.m.) across paired samples; signal-to-noise ratios (SNR) are reported as |mean difference|/s.e.m. of the difference.

### Convergence and robustness checks

Numerically exact methods (sparse matrix exponentials for the eight-site model and the exact singlet-yield expression for the unified Hamiltonians) were used throughout because the Hilbert spaces involved (≤ 2 dimensions) are small; tensor-network methods would be required for the full chain. The exact singlet-yield expression removes time-step and integration-window errors; finite-window convergence is shown in **Supplementary Fig. 4c** (yields within 0.3% of their infinite-time values at 6 µs). For the eight-site model, the effect of the unknown hop rates was assessed by repeating every calculation with the two bracketing values.

### Summary of spin Hamiltonians

(1) Structural coupling map (Fig. 1c,d): pairwise J(r) and d(r) for the eight main-chain spins; no dynamics. (2) Eight-site coherence-propagation model (Fig. 2a–c): Zeeman, FMN hyperfine, and all-pairs exchange and dipolar coupling, with dephasing and unital electron-hop dissipators; no O site and no recombination. (3) Unified O –FMN–N3 radical-pair model (Figs 2d, 3a,b, Supplementary Fig. 4 and Supplementary Figs 1 and 2a): Zeeman, FMN hyperfine, O –FMN exchange and axial dipolar coupling, Haberkorn recombination (kS = kT = 10 s ¹ unless stated), FMN–N3 exchange and dipolar coupling, N3 dephasing and the FMN→N3 hop channel. (4) Extended unified model (Figs 3c,d and Supplementary Fig. 2b): as (3), extended to N1b and N4 with the corresponding couplings, dephasing and hop channels. (5) Charge-resolved O –FMN–N3 model (Fig. 4 and Supplementary Fig. 3): three redox configurations with spin-conserving, detailed-balance FMN N3 transfer, N3 re-oxidation by N1b, isotropic Fe–S relaxation and Haberkorn recombination in the FMN-semiquinone configurations. (6) Unital spin-hop control (Fig. 4c): as (3) with each flip-flop jump operator supplemented by its reverse.

### Software

Simulations were performed in Python 3.13 with QuTiP 5.3.1, NumPy 2.5 and SciPy 1.18 (sparse matrix exponentials and k-d tree queries). Structures were parsed with Biopython 1.88, figures were prepared with Matplotlib 3.11, and dipolar tensors were verified against RadicalPy 1.0.9.

## Code availability

All code used in this study, including the structure-processing pipeline, the eight-site coherence-propagation model, the unified radical-pair Hamiltonians and the exact singlet-yield solver, is available at Supplementary material.

## Data availability

All numerical results underlying the figures and supplementary tables are provided as Source Data with this paper. Structural coordinates were obtained from the Protein Data Bank (PDB entry: 5XTD).

## AI Use Declaration

AI tools were used only for English grammar correction and language polishing.

## Authors’ contributions

**Ji-Yong Sung** and **Jae-Ho Cheong** conceived and designed the study. **Ji-Yong Sung** developed the theoretical framework and computational models, performed the quantum spin-dynamics simulations, analysed the data and prepared the figures. Both authors interpreted the results and discussed their physical and biological implications. **Ji-Yong Sung** wrote the original manuscript. **Jae-Ho Cheong** acquired funding. Both authors supervised the research, reviewed and edited the manuscript, and approved the final version.

## Acknowledgements

N/A

## Reference

1. Kussmaul, L. and J. Hirst, The mechanism of superoxide production by NADH:ubiquinone oxidoreductase (complex I) from bovine heart mitochondria. Proceedings of the National Academy of Sciences, 2006. 103(20): p. 7607–7612.

2. Hirst, J., Towards the molecular mechanism of respiratory complex I. Biochemical Journal, 2009. 425(2): p. 327–339.

3. Bazil, J.N., et al., Determining the origins of superoxide and hydrogen peroxide in the mammalian NADH:ubiquinone oxidoreductase. Free Radical Biology and Medicine, 2014. 77: p. 121–129.

4. Chouchani, E.T., et al., Ischaemic accumulation of succinate controls reperfusion injury through mitochondrial ROS. Nature, 2014. 515(7527): p. 431–435.

5. Schapira, A.H.V., et al., Mitochondrial Complex I Deficiency in Parkinson’s Disease. Journal of Neurochemistry, 1990. 54(3): p. 823–827.

6. Sung, J.-Y., L. M. A. & Cheong, J.-H., Structure constrained spin responses of an FMN superoxide radical-pair model in mitochondrial complex I. https://www.biorxiv.org/content/10.64898/2026.07.15.738821v3, 2026.

7. Rodgers, C.T. and P.J. Hore, Chemical magnetoreception in birds: The radical pair mechanism. Proceedings of the National Academy of Sciences, 2009. 106(2): p. 353–360.

8. Zadeh-Haghighi, H. and C. Simon, Magnetic field effects in biology from the perspective of the radical pair mechanism. Journal of The Royal Society Interface, 2022. 19(193).

9. Evans, E.W., et al., Magnetic field effects in flavoproteins and related systems. Interface Focus, 2013. 3(5).

10. Hinchliffe, P. and L.A. Sazanov, Organization of Iron-Sulfur Clusters in Respiratory Complex I. Science, 2005. 309(5735): p. 771–774.

11. Couch, V.A., E.S. Medvedev, and A.A. Stuchebrukhov, Electrostatics of the FeS clusters in respiratory complex I. Biochimica et Biophysica Acta (BBA) - Bioenergetics, 2009. 1787(10): p. 1266–1271.

12. Gnandt, E., et al., The multitude of iron–sulfur clusters in respiratory complex I. Biochimica et Biophysica Acta (BBA) - Bioenergetics, 2016. 1857(8): p. 1068–1072.

13. Harvey, S.M. and M.R. Wasielewski, Photogenerated Spin-Correlated Radical Pairs: From Photosynthetic Energy Transduction to Quantum Information Science. Journal of the American Chemical Society, 2021. 143(38): p. 15508–15529.

14. Hore, P.J., Spin chemistry in living systems. National Science Review, 2024. 11(9).

15. Adams, B., I. Sinayskiy, and F. Petruccione, An open quantum system approach to the radical pair mechanism. Scientific Reports, 2018. 8(1).

16. Denton, M.C.J., et al., Magnetosensitivity of tightly bound radical pairs in cryptochrome is enabled by the quantum Zeno effect. Nature Communications, 2024. 15(1).

17. Kattnig, D.R. and P.J. Hore, The sensitivity of a radical pair compass magnetoreceptor can be significantly amplified by radical scavengers. Scientific Reports, 2017. 7(1).

18. Kattnig, D.R., Radical-Pair-Based Magnetoreception Amplified by Radical Scavenging: Resilience to Spin Relaxation. The Journal of Physical Chemistry B, 2017. 121(44): p. 10215–10227.

19. Babcock, N.S. and D.R. Kattnig, Electron–Electron Dipolar Interaction Poses a Challenge to the Radical Pair Mechanism of Magnetoreception. The Journal of Physical Chemistry Letters, 2020. 11(7): p. 2414–2421.

20. Verkhovskaya, M.L., et al., Real-time electron transfer in respiratory complex I. Proceedings of the National Academy of Sciences, 2008. 105(10): p. 3763–3767.

21. Guo, R., et al., Architecture of Human Mitochondrial Respiratory Megacomplex I2III2IV2. Cell, 2017. 170(6): p. 1247–1257.e12.

22. Le Breton, N., et al., Using Hyperfine Electron Paramagnetic Resonance Spectroscopy to Define the Proton-Coupled Electron Transfer Reaction at Fe–S Cluster N2 in Respiratory Complex I. Journal of the American Chemical Society, 2017. 139(45): p. 16319–16326.

23. Cock, P.J.A., et al., Biopython: freely available Python tools for computational molecular biology and bioinformatics. Bioinformatics, 2009. 25(11): p. 1422–1423.

24. Ohnishi, T. and E. Nakamaru-Ogiso, Were there any “misassignments” among iron–sulfur clusters N4, N5 and N6b in NADH-quinone oxidoreductase (complex I)? Biochimica et Biophysica Acta (BBA) - Bioenergetics, 2008. 1777(7-8): p. 703–710.

25. Moser, C.C., et al., Electron tunneling chains of mitochondria. Biochimica et Biophysica Acta (BBA) - Bioenergetics, 2006. 1757(9-10): p. 1096–1109.

26. Hirst, J. and M.M. Roessler, Energy conversion, redox catalysis and generation of reactive oxygen species by respiratory complex I. Biochimica et Biophysica Acta (BBA) - Bioenergetics, 2016. 1857(7): p. 872–883.

27. Hirst, J., Mitochondrial Complex I. Annual Review of Biochemistry, 2013. 82(1): p. 551–575.

28. Moser, C.C., et al., Nature of biological electron transfer. Nature, 1992. 355(6363): p. 796–802.

29. Antill, L.M. and E. Vatai, RadicalPy: A Tool for Spin Dynamics Simulations. Journal of Chemical Theory and Computation, 2024. 20(21): p. 9488–9499.

30. Bridges, H.R., E. Bill, and J. Hirst, Mössbauer Spectroscopy on Respiratory Complex I: The Iron–Sulfur Cluster Ensemble in the NADH-Reduced Enzyme Is Partially Oxidized. Biochemistry, 2011. 51(1): p. 149–158.

31. de Vries, S., et al., Electron Tunneling Rates in Respiratory Complex I Are Tuned for Efficient Energy Conversion. Angewandte Chemie International Edition, 2015. 54(9): p. 2844–2848.

32. Marques, H.M., Electron transfer in biological systems. JBIC Journal of Biological Inorganic Chemistry, 2024. 29(7-8): p. 641–683.

33. Hayashi, T. and A.A. Stuchebrukhov, Quantum Electron Tunneling in Respiratory Complex I. The Journal of Physical Chemistry B, 2011. 115(18): p. 5354–5364.

34. Hayashi, T. and A. Stuchebrukhov, Electron tunneling pathways in respiratory complex I: The role of the internal water between the enzyme subunits. Journal of Electroanalytical Chemistry, 2011. 660(2): p. 356–359.

35. Fiedorczuk, K., et al., Atomic structure of the entire mammalian mitochondrial complex I. Nature, 2016. 538(7625): p. 406–410.

36. Steiner, U.E. and T. Ulrich, Magnetic field effects in chemical kinetics and related phenomena. Chemical Reviews, 2002. 89(1): p. 51–147.

37. Gorini, V., A. Kossakowski, and E.C.G. Sudarshan, Completely positive dynamical semigroups of N-level systems. Journal of Mathematical Physics, 1976. 17(5): p. 821–825.

38. Plenio, M.B. and S.F. Huelga, Dephasing-assisted transport: quantum networks and biomolecules. New Journal of Physics, 2008. 10(11).

39. Hore, P.J. and H. Mouritsen, The Radical-Pair Mechanism of Magnetoreception. Annual Review of Biophysics, 2016. 45(1): p. 299–344.

40. Page, C.C., et al., Natural engineering principles of electron tunnelling in biological oxidation– reduction. Nature, 1999. 402(6757): p. 47–52.

41. Haberkorn, R., Density matrix description of spin-selective radical pair reactions. Molecular Physics, 2006. 32(5): p. 1491–1493.

42. Ivanov, K.L., et al., Consistent Treatment of Spin-Selective Recombination of a Radical Pair Confirms the Haberkorn Approach. The Journal of Physical Chemistry A, 2010. 114(35): p. 9447–9455.

43. Jones, J.A. and P.J. Hore, Spin-selective reactions of radical pairs act as quantum measurements. Chemical Physics Letters, 2010. 488(1-3): p. 90–93.

44. Fay, T.P., L.P. Lindoy, and D.E. Manolopoulos, Spin-selective electron transfer reactions of radical pairs: Beyond the Haberkorn master equation. The Journal of Chemical Physics, 2018. 149(6).

45. Hayashi, T. and A.A. Stuchebrukhov, Electron tunneling in respiratory complex I. Proceedings of the National Academy of Sciences, 2010. 107(45): p. 19157–19162.

46. Al-Mohy, A.H. and N.J. Higham, Computing the Action of the Matrix Exponential, with an Application to Exponential Integrators. SIAM Journal on Scientific Computing, 2011. 33(2): p. 488–511.

47. Johansson, J.R., P.D. Nation, and F. Nori, QuTiP: An open-source Python framework for the dynamics of open quantum systems. Computer Physics Communications, 2012. 183(8): p. 1760–1772.

