## supplementary information for "Kinetic gating by an iron–sulfur chain modulates flavin radical-pair spin chemistry in mitochondrial Complex I"

**Contents**

Supplementary Note 1. Coherence generated by one-directional electron-hop operators in the control trajectory

Supplementary Note 2. Origin of the difference from the near-contact maximum reported previously

Supplementary Note 3. Robustness to the hyperfine coupling constant

Supplementary Note 4. Charge-resolved model: construction and checks

Supplementary Note 5. Spin-hop model: sensitivity, recombination-rate dependence and convergence

Supplementary Note 6. Robustness of the charge-resolved results

Supplementary Fig. 1. Network effect on the singlet yield as a function of the recombination rates (spin-hop model)

Supplementary Fig. 2. Distance dependence at fast recombination and the N3→N1b sweep at *r* = 6.5 Å (spin-hop model)

Supplementary Fig. 3. Robustness of the charge-resolved model

Supplementary Fig. 4. Spin-hop model: sensitivity, FMN→N3 rate and integration window

Supplementary Fig. 5. One-directional versus unital hop operators in the eight-site coherence-propagation model

Supplementary Table 1. Geometry, coupling constants and electron-transfer rates of the main-chain links

Supplementary Table 2. Location of the composite-score maximum under combinations of model choices

Supplementary Table 3. Key results with the anionic (1.233 mT) and neutral (1.108 mT) flavin hyperfine fields

Supplementary Table 4. Singlet yields for the model comparison in Fig. 4c with the two exchange conventions

Supplementary Table 5. Charge-resolved singlet yields as a function of the FMN→N3 driving force

Supplementary Table 6. Recombination-rate grid of the charge-resolved network effect at *r* = 3.3 Å (Fig. 4d)

**Supplementary Note 1. Coherence generated by one-directional electron-hop operators in the control trajectory**

In the unified radical-pair models (Figs 2d and 3), electron transfer between adjacent centres *i* and *j* is represented by the one-directional jump operator *L* = √*k*_ET_ σ̂^+^_i_σ̂^−^_j_. Because this operator is not normal (*LL*^†^ ≠ *L*^†^*L*), the corresponding Lindblad dissipator does not leave the identity unchanged:

𝒟[*L*](𝟙) = *LL*^†^ − *L*^†^*L* = *k*_ET_(*P*^↑^_i_*P*^↓^_j_ − *P*^↓^_i_*P*^↑^_j_) = *k*_ET_(*Ŝ*_z,i_ − *Ŝ*_z,j_),

where *P*^↑^ and *P*^↓^ are the single-spin projectors. The maximally mixed state is therefore not stationary: each hop channel builds up longitudinal polarization of opposite sign on the two centres it connects. On FMN, this polarization precesses about the local field, which is dominated by the randomly oriented semiclassical hyperfine field (*a*_HF_ = 1.233 mT) rather than by the 0.05 mT static field along *z*, and so acquires a transverse component. A control trajectory in which every centre starts in 𝟙/2 therefore develops a nonzero ⟨*Ŝ*^+^_FMN_⟩ without any injected coherence.

Adding the reverse operator, *L*′ = √*k*_ET_ σ̂^+^_j_σ̂^−^_i_, at the same rate removes this term, because 𝒟[*L*′](𝟙) = *k*_ET_(*Ŝ*_z,j_ − *Ŝ*_z,i_) cancels 𝒟[*L*](𝟙). The combined dissipator is then unital, the maximally mixed state is stationary and no polarization is generated. The eight-site coherence-propagation model of Fig. 2a–c therefore uses these unital hops; the one-directional operator is used there only for comparison (Supplementary Fig. 5).

Numerically, applying the Liouvillian of the eight-site model to the vectorised maximally mixed state, 𝟙/2^8^, gives a vector of norm 3.8 × 10^8^ s^−1^ with one-directional hop channels, ≈ 10^−9^ s^−1^ with unital hops and exactly zero when the hop channels are removed. With one-directional hops, propagating the control state for two of the twelve hyperfine orientations used in Fig. 2 (the first two of the fixed-seed set) gave |⟨*Ŝ*^+^_FMN_⟩| = 0.051 and 0.096 at 5 ns, increasing monotonically over the window; with unital hops, or without hop channels, no coherence appeared (Supplementary Fig. 5c). These values are comparable with the baseline-subtracted coherence remaining at FMN in the one-directional excited trajectory (0.12 at 5 ns; Supplementary Fig. 5a), so with one-directional operators the control signal must be removed before coherence propagation can be assessed.

Because the master equation is linear, the difference between the excited and control trajectories obeys the same equation as each trajectory. With the excited site *k* prepared in |+*x*⟩ and all other sites in 𝟙/2, the initial difference is |+*x*⟩⟨+*x*| ⊗ (𝟙/2)^⊗7^ − (𝟙/2)^⊗8^ = *Ŝ*_x,k_ ⊗ (𝟙/2)^⊗7^. Propagating this traceless operator directly gives the baseline-subtracted signal exactly, without statistical cancellation between two large trajectories. All traces in Fig. 2a–c and Supplementary Fig. 5a,b were obtained in this way; with unital hops the subtraction is a precaution, because the control trajectory is stationary.

The spurious polarization is a consequence of representing electron transfer as a one-directional spin flip-flop between permanently paramagnetic sites, rather than as a change in site occupancy. In the unified radical-pair models (Figs 2d and 3), singlet yields are evaluated from a triplet-born O₂–FMN pair rather than from the difference of two trajectories, so no baseline subtraction is involved. The same polarizing drive nevertheless acts on FMN in those models and underlies the suppression of the singlet yield that they predict at slow recombination (Fig. 4c and Supplementary Note 4); it is absent from the charge-resolved model (Fig. 4).

**Supplementary Note 2. Origin of the difference from the near-contact maximum reported previously**

Our earlier analysis of the isolated FMN–O₂ pair placed the maximum of the composite score *H*(*r*) near 3.3 Å (ref. 6), whereas the network-OFF calculation in Fig. 3a places it at 6.0–6.5 Å. To identify the cause, we re-implemented the two-spin (network-OFF) model as a vectorised solver that reproduces the singlet yields of the unified Hamiltonian used in this work to machine precision (largest difference 4 × 10^−16^), and switched six model choices independently between the two analyses:

**Hyperfine field.** This work: single semiclassical field of fixed magnitude 1.233 mT and random orientation. Earlier analysis: Schulten–Wolynes field from four ^14^N and twelve ^1^H nuclei (RadicalPy dataset flavin_anion; root-sum-square 1.233 mT).

**O₂–FMN dipolar term.** This work: axial point-dipole term included. Earlier analysis: not included (exchange only).

**Steric test.** This work: no protein or non-FMN heteroatom within 3.0 Å and no FMN atom within 2.8 Å. Earlier analysis: distance to every non-FMN heavy atom greater than its van der Waals radius plus a 1.52 Å probe; FMN atoms not tested.

**Field sensitivity *M*_B_.** This work: finite-difference derivative at 0.05 ± 0.02 mT. Earlier analysis: |*Y*_S_(50 mT) − *Y*_S_(0.05 mT)|.

**Dephasing sensitivity *C*_γ_.** This work: finite difference between 1.5 × 10^7^ and 6 × 10^7^ s^−1^. Earlier analysis: |*Y*_S_(γ_low_) − *Y*_S_(γ_high_)|, taken here as 10^6^ and 10^9^ s^−1^.

**Normalisation of *M*_B_ and *C*_γ_.** This work: divided by the maximum. Earlier analysis: min–max scaled to 0–1.

Van der Waals radii were C 1.70, N 1.55, O 1.52, S 1.80, P 1.80 and Fe 1.94 Å. Accessibility was evaluated from 1.5 × 10^6^ points sampled uniformly in *r* between 2.45 and 9.05 Å around the isoalloxazine centroid of PDB 5XTD and expressed as the accepted fraction in 0.1 Å bins. Spin responses were computed on the same 0.1 Å grid (*r* = 2.5–9.0 Å) with 400 paired hyperfine samples, *k*_S_ = *k*_T_ = 10^6^ s^−1^ and *k*_STD_ = 3 × 10^7^ s^−1^. The values of γ_low_ and γ_high_ were not stated in the earlier report; the choice made here reproduces its dephasing-response profile, which is largest near contact and approaches a plateau beyond about 4–5 Å.

With all six earlier choices, *H*(*r*) peaked at 3.4 Å, reproducing the earlier result (3.25 Å). With all six choices of this work it peaked at 5.8 Å, consistent with Fig. 3a given the coarser grid and larger sample used there. Supplementary Table 2 lists the peak location when one choice at a time is switched. Starting from the earlier configuration, only the steric test moved the maximum away from contact: excluding positions that overlap FMN moved it to 5.9 Å with the van der Waals criterion and to 4.1 Å with the criterion of this work, whereas each of the other five changes left it between 2.5 and 3.8 Å. Starting from the configuration of this work, no single change restored a near-contact maximum (5.2–9.0 Å).

The reason is evident from the accessible fractions. At *r* = 3.3 Å, 36% of candidate positions pass the van der Waals test when only protein and other non-FMN atoms are considered, but only 0.3% remain when FMN atoms are also tested (39% and 1.9%, respectively, with the criterion of this work). Most positions counted as accessible near contact in the earlier analysis therefore overlap the flavin itself. Because the dephasing response is largest at short separation, these positions dominate the earlier composite score; once they are excluded, the maximum moves to the intermediate separations where both accessibility and spin sensitivity are appreciable.

**Supplementary Note 3. Robustness to the hyperfine coupling constant**

The reference hyperfine coupling, *a*_HF_ = 1.233 mT, carried forward from our earlier work is the root-sum-square isotropic hyperfine field, [Σ_k_ *a*_k_^2^*I*_k_(*I*_k_ + 1)]^1/2^, of the sixteen-nucleus flavin dataset distributed with RadicalPy as flavin_anion (four ^14^N and twelve ^1^H), which describes the anionic FMN^•−^ radical. The corresponding dataset for the neutral FMNH^•^ semiquinone (flavin_neutral; four ^14^N and seventeen ^1^H) gives 1.108 mT. Because the radical pair is treated here as FMNH^•^–O₂^•−^, we repeated the key calculations with *a*_HF_ = 1.108 mT, leaving every other parameter and the hyperfine orientations unchanged.

No conclusion changed (Supplementary Table 3). The network-induced change in singlet yield at 3.3 Å differed by 1.5%, the chain-truncation steps by less than 0.0002, and the midpoints of the kinetic threshold by less than 0.1%. All 25 entries of the *k*_S_–*k*_T_ grid of Supplementary Fig. 1 kept their sign; the largest change in a moderate entry was 2.9 percentage points (*k*_S_ = 10^8^, *k*_T_ = 10^5^ s^−1^: +18.6% and +21.5%). The distance dependence at fast recombination was also preserved (+8.5% and +9.3% at 3.3 Å; −2.1% and −2.0% at 6.5 Å). With FMN atoms excluded from the steric test, the network-OFF score again peaked at intermediate separation (6.5 Å with 1.108 mT, 6.0 Å with 1.233 mT in this 1,000-sample check) and the network-ON score rose to a broad maximum beyond 6.5 Å in both cases, without a near-contact maximum. The dephasing sensitivity remained lower with network coupling at every separation (ratio ON/OFF 0.50–0.91 with 1.108 mT; 0.45–0.93 with 1.233 mT).

The location of the network-OFF maximum varies by about 0.5 Å between Monte Carlo samples because the network-OFF field sensitivity is small at intermediate separations (relative standard error 17–25% at 6–7 Å with 2,000 samples). This uncertainty does not affect the absence of a near-contact maximum.

**Supplementary Note 4. Charge-resolved model: construction and checks**

The charge-resolved model is described in Methods (Fig. 4a). Three checks were performed. (i) With FMN→N3 transfer switched off and the system started in configuration c2, singlet yields agree with those of the QuTiP implementation of the isolated pair (radical_pair.py) to within 2 × 10^−5^ for every hyperfine orientation tested (tests/test_charge_resolved.py). (ii) Singlet, triplet and escape yields sum to one to within 10^−6^. (iii) The isotropic relaxation operators give *T*₁^−1^ = *T*₂^−1^ equal to the nominal rate to within 10^−9^ (relative). A NumPy implementation of the spin-hop model (numpy_models.py) reproduces the QuTiP results to within 2 × 10^−5^; its default hop direction matches the operator √k σ̂^+^(FMN) σ̂^−^(N3) of radical_pair.py, and reversing the operator direction leaves orientation-averaged yields unchanged within sampling error.

**Why the spin-hop operator suppresses the singlet yield.** The jump operator √k σ̂^+^_i_σ̂^−^_j_ is non-unital: it transfers population in one direction only and, combined with fast dephasing of the Fe–S spin, drives the FMN spin towards a polarized state. Adding the reverse operator at the same rate makes the dissipator unital and removes this polarizing drive; the singlet yield then moves towards the incoherent limit, as in the charge-resolved model (Fig. 4c and Supplementary Table 4).

**Supplementary Note 5. Spin-hop model: sensitivity, recombination-rate dependence and convergence**

*All results in this Note refer to the phenomenological spin-hop model (Figs 2d and 3).*

Network coupling increased the combined spin sensitivity at every separation at which it could be resolved, by 2–3-fold at r ≤ 4 Å and by 7–35-fold at 4.5–7 Å (Supplementary Fig. 4a). This increase arises entirely from the magnetic-field term: the dephasing sensitivity, *C*_γ_, was lower with network coupling at every separation (by 6–57%; by 35% at 3.3 Å).

The direction of the nearest-neighbour effect also depends on the recombination kinetics (Supplementary Fig. 1). Exact yields on a *k*_S_–*k*_T_ grid at *r* = 3.3 Å showed that coupling to N3 suppressed *Y*_S_ only when recombination was slow (*k*_S_ = *k*_T_ = 10^5^ or 10^6^ s^−1^, and *k*_S_ = 10^5^, *k*_T_ = 10^6^ s^−1^). Whenever either rate was 10^7^ s^−1^ or faster, coupling increased *Y*_S_, by 5–23% for most combinations and by about 80% to more than sevenfold when *k*_T_ ≥ 10^8^ s^−1^, where *Y*_S_ itself is small. At the diffusion-influenced rate estimated for a flavin semiquinone–superoxide pair (*k*_S_ ≈ 4 × 10^8^ s^−1^) with *k*_T_ = 10^6^ s^−1^, coupling increased *Y*_S_ by 8.5%. This increase is confined to short separations: at these rates the N3 contribution decreases with distance, changes sign near 5.5 Å and lowers *Y*_S_ by 1.9% at 6.5 Å, where oxygen is sterically accessible (Supplementary Fig. 2a). With both rates fast (*k*_S_ = *k*_T_ = 10^8^ s^−1^), coupling increased *Y*_S_ by 46–82% at every separation from 2.6 to 9.0 Å. Repeating the N3→N1b rate sweep at *r* = 6.5 Å gave the same threshold, with similar midpoints (1.3 × 10^6^ and 7.3 × 10^6^ s^−1^), at the two slower reaction rates; at *k*_S_ = *k*_T_ = 10^8^ s^−1^, however, the N1b contribution peaked near 10^8^ s^−1^ and became negative at faster hopping (Supplementary Fig. 2b). The absence of a resonance maximum in Fig. 3d therefore holds at slow recombination but not at every separation when recombination is fast. All values reported here use the exact singlet-yield expression (Methods).

By contrast, the singlet yield was insensitive to the FMN→N3 rate, which is several orders of magnitude faster than the reaction rate (Supplementary Fig. 4b). Across 10^8^–10^11^ s^−1^, *Y*_S_ with network coupling varied by 2.4%, and by less than 0.8% within fourfold of the Moser–Dutton estimate (8.5 × 10^9^ s^−1^); network suppression persisted across the entire range.

Finally, we confirmed that the reported singlet yields are not sensitive to the finite duration of the integration window (Supplementary Fig. 4c). Both conditions, computed from the same underlying Hamiltonian, deviated from their infinite-time yields by 2.0% (OFF) and 1.4% (ON) at 4 µs and by less than 0.3% at 6 µs, the window used for our time-domain cross-checks (Methods); at these rates the radical pair has therefore essentially fully reacted within 6 µs. This is shorter than the slowest hop included in the extended Hamiltonians (N1b→N4, τ ≈ 16 µs) and more than twentyfold shorter than the chain's slowest electron-hop step (N5→N6a, τ ≈ 137 µs). None of the results presented in this study therefore probe genuine microsecond-to-millisecond multi-turnover dynamics; all reported effects are established on the timescale of a single radical-pair reaction event.

**Supplementary Note 6. Robustness of the charge-resolved results**

**Exchange convention.** The unified spin-hop model (Figs 2d and 3) writes the O₂–FMN exchange as +J Ŝ_O₂_·Ŝ_FMN_, whereas the charge-resolved and comparison calculations (Fig. 4) use −2J Ŝ_O₂_·Ŝ_FMN_ with the same J(r). Supplementary Table 4 repeats the Fig. 4c comparison with both conventions (400 paired orientations). The largest difference, 0.011, occurs for the isolated pair at *r* = 3.3 Å and *k*_S_ = 4 × 10^8^ s^−1^; no sign or conclusion changes. The difference between the spin-hop yields quoted in the text (0.154 in Fig. 2d and 0.153 in Fig. 4c) arises from this convention.

**FMN→N3 driving force.** Because the one-electron potentials of the Complex I flavin are uncertain, the charge-resolved calculations were repeated with ΔG = −0.05, −0.09, −0.164 and −0.20 eV, with *k*_f_ from the Moser–Dutton relation and *k*_b_ from detailed balance at 310 K (Supplementary Table 5; 150 paired orientations). Reduced-chain yields, with N3 held reduced or re-oxidizable by N1b, changed by at most 3 × 10^−4^, and the N3→N1b rate sweep at *r* = 3.3 Å and *k*_S_ = *k*_T_ = 10^6^ s^−1^ was identical for ΔG = −0.09 and −0.164 eV. Once N3 has been re-oxidized, reversible FMN⇄N3 shuttling is much faster than recombination for every ΔG, so the spin correlation is randomized irrespective of *k*_f_ and *k*_b_. Yields starting from an oxidized N3, with onward transfer from N3 to N1b, depend strongly on ΔG and on the recombination rates.

**Fe–S dephasing operator.** In the unified spin-hop model the Fe–S dephasing operator √γ σ̂z gives a transverse decay rate of 2γ, twice that of the eight-site model. With √(γ/2) σ̂z instead, Δ*Y*_S_ changed from −0.0123 to −0.0142 at 3.3 Å and from −0.0104 to −0.0197 at 6.5 Å (200 paired orientations, *k*_S_ = *k*_T_ = 10^6^ s^−1^), without a change of sign.

**Sign of the charge-resolved effect at *k*_T_ > *k*_S_.** In the 5 × 5 grid of Fig. 4d (Supplementary Table 6), the isolated-pair yield lay below the incoherent limit at all ten points with *k*_T_ > *k*_S_, yet the network lowered *Y*_S_ at nine of them (by 1–13%), moving it away from the limit. The tendency towards the incoherent limit is therefore restricted to *k*_S_ ≥ *k*_T_.

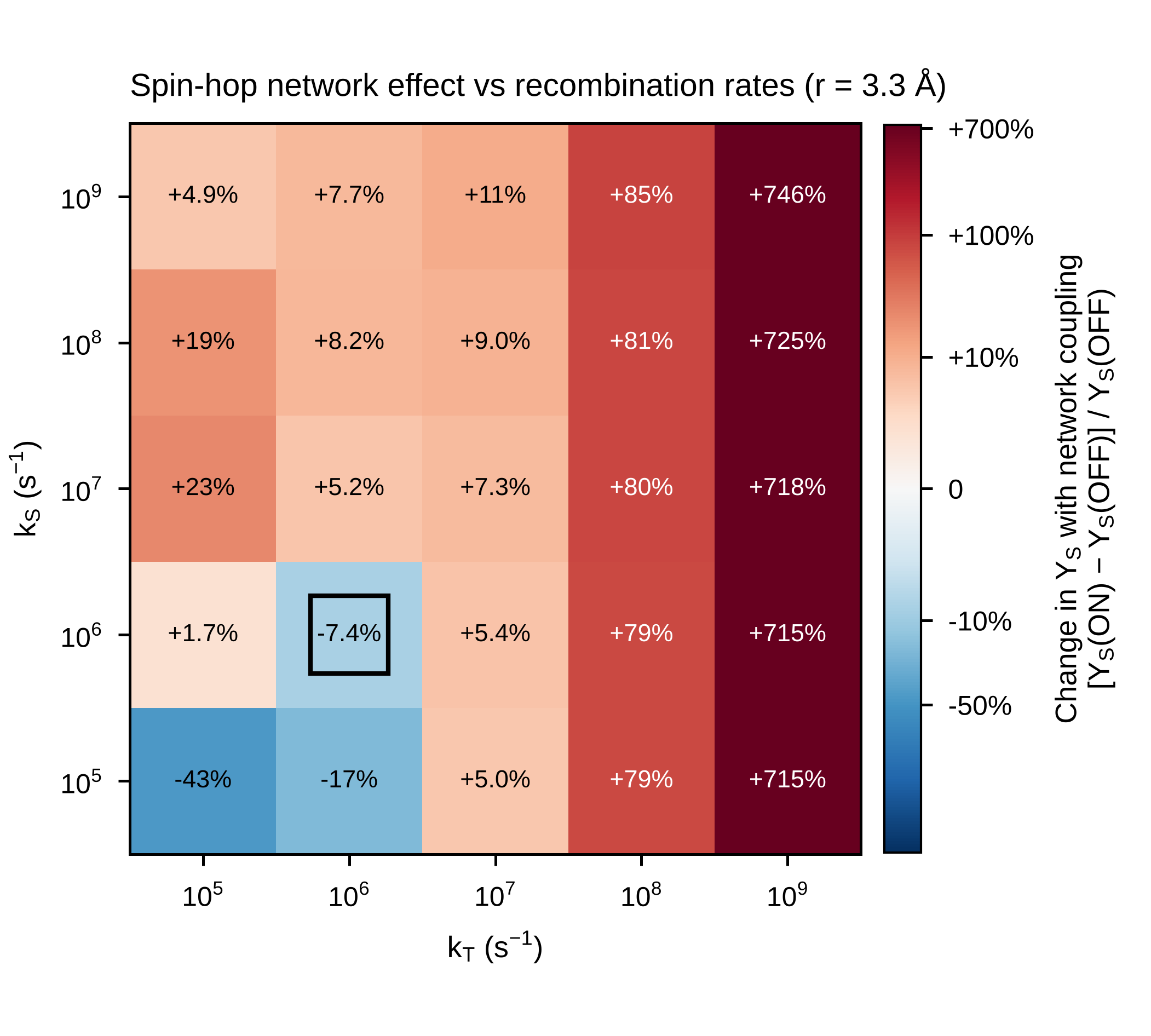

**Supplementary Fig. 1 | Network effect on the singlet yield as a function of the recombination rates (spin-hop model).** Relative change in the singlet yield on coupling FMN to N3, [*Y*_S_(ON) − *Y*_S_(OFF)]/*Y*_S_(OFF), at *r* = 3.3 Å on a 5 × 5 grid of singlet and triplet recombination rates (10^5^–10^9^ s^−1^). Yields were computed from the unified O₂–FMN–N3 spin-hop Hamiltonian, evaluated exactly at infinite time and averaged over 400 paired hyperfine orientations. The box marks the baseline rates, *k*_S_ = *k*_T_ = 10^6^ s^−1^. The colour scale is non-linear. Coupling lowers the yield only when both rates are slow; for the distance dependence at fast recombination, see Supplementary Fig. 2a.

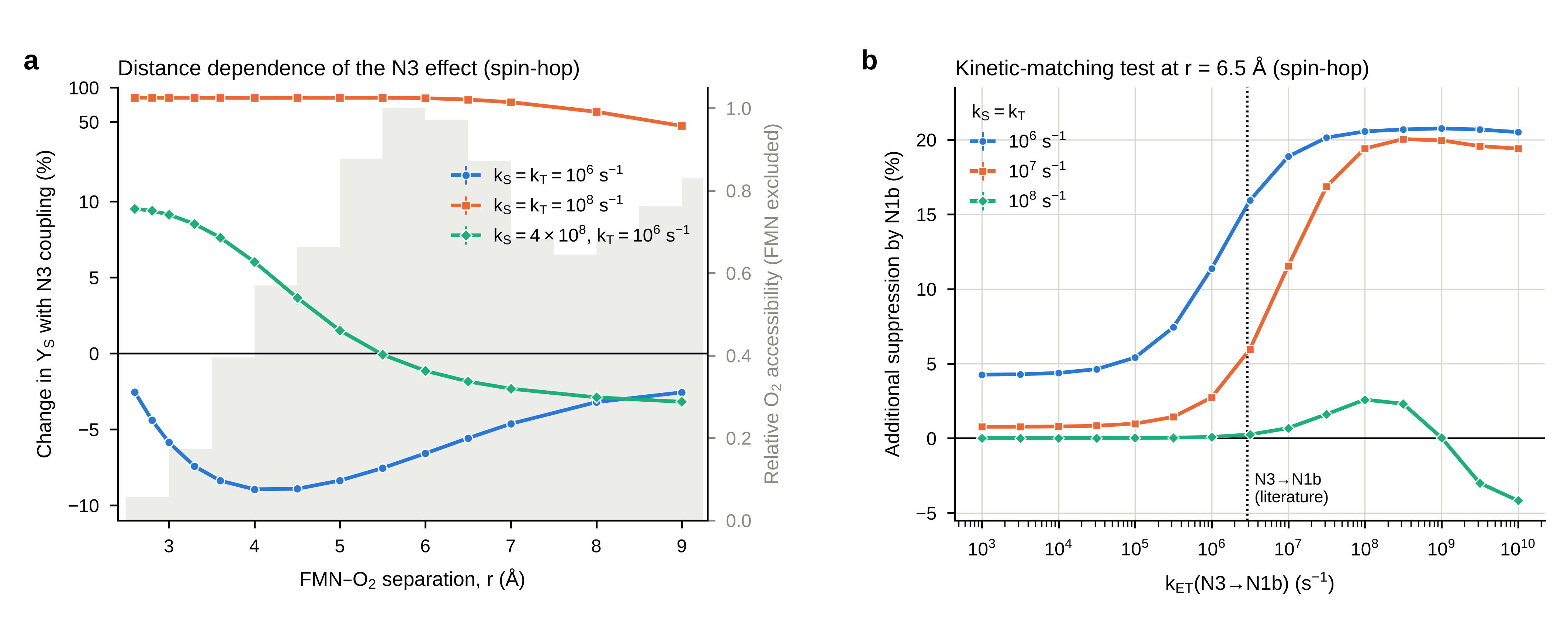

**Supplementary Fig. 2 | Distance dependence of the network effect at fast recombination, and the N3→N1b rate sweep at an accessible separation (spin-hop model).** **a**, Relative change in the singlet yield on coupling FMN to N3, as a function of FMN–O₂ separation, for slow recombination (*k*_S_ = *k*_T_ = 10^6^ s^−1^, blue), fast recombination in both channels (*k*_S_ = *k*_T_ = 10^8^ s^−1^, orange) and a fast singlet channel alone (*k*_S_ = 4 × 10^8^, *k*_T_ = 10^6^ s^−1^, green). The vertical axis is linear within ±10% and logarithmic beyond. Grey, relative O₂ accessibility with FMN atoms excluded (as in Fig. 3a). With a fast singlet channel alone, the effect changes sign near 5.5 Å. 1,000 paired hyperfine orientations. **b**, Additional suppression of the singlet yield contributed by N1b, [*Y*_S_(+N3) − *Y*_S_(+N3+N1b)]/*Y*_S_(+N3), as a function of the N3→N1b hop rate at *r* = 6.5 Å (compare Fig. 3d, *r* = 3.3 Å). Dotted line, literature N3→N1b rate (2.9 × 10^6^ s^−1^). At *k*_S_ = *k*_T_ = 10^8^ s^−1^ the contribution peaks near 10^8^ s^−1^ and becomes negative at faster hopping. 400 paired hyperfine orientations. In both panels, yields were evaluated exactly at infinite time; error bars, s.e.m.

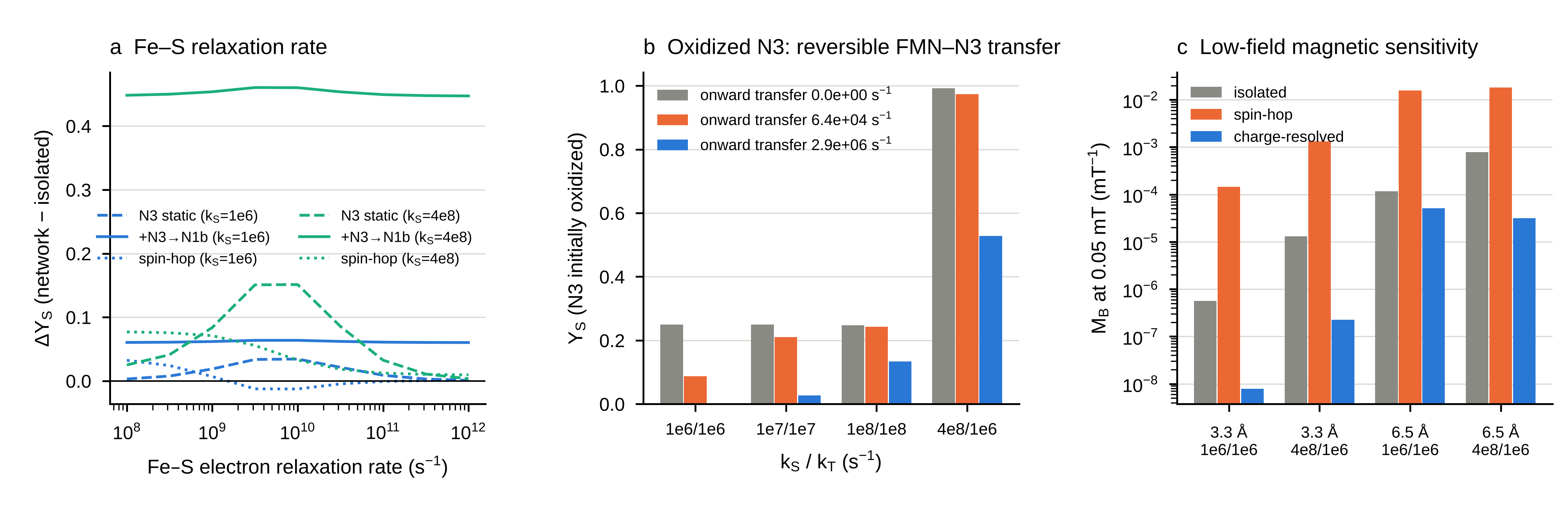

**Supplementary Fig. 3 | Robustness of the charge-resolved model.** **a**, Network-induced change in the singlet yield at *r* = 3.3 Å as a function of the Fe–S electron relaxation rate, for the charge-resolved model with N3 held reduced (dashed) or re-oxidized by N1b at the literature rate (solid), and for the spin-hop model (dotted), at *k*_S_ = *k*_T_ = 10^6^ s^−1^ (blue) and *k*_S_ = 4 × 10^8^, *k*_T_ = 10^6^ s^−1^ (green). The gated contribution is nearly independent of the relaxation rate, whereas that of a static reduced N3 peaks near 10^10^ s^−1^ and vanishes in the motional-narrowing limit. **b**, Singlet yield when N3 is initially oxidized, with reversible FMN–N3 transfer (detailed balance) and onward transfer from N3 at the rates shown. Fast onward transfer quenches the radical pair. **c**, Low-field magnetic sensitivity, *M*_B_ = |⟨*Y*_S_(0.07 mT) − *Y*_S_(0.03 mT)⟩|/0.04 mT, for the isolated pair, the spin-hop model and the charge-resolved model, without additional O₂•^−^ relaxation. The spin-hop network increases *M*_B_ by one to two orders of magnitude; the charge-resolved network lowers it. 150 paired hyperfine orientations per point.

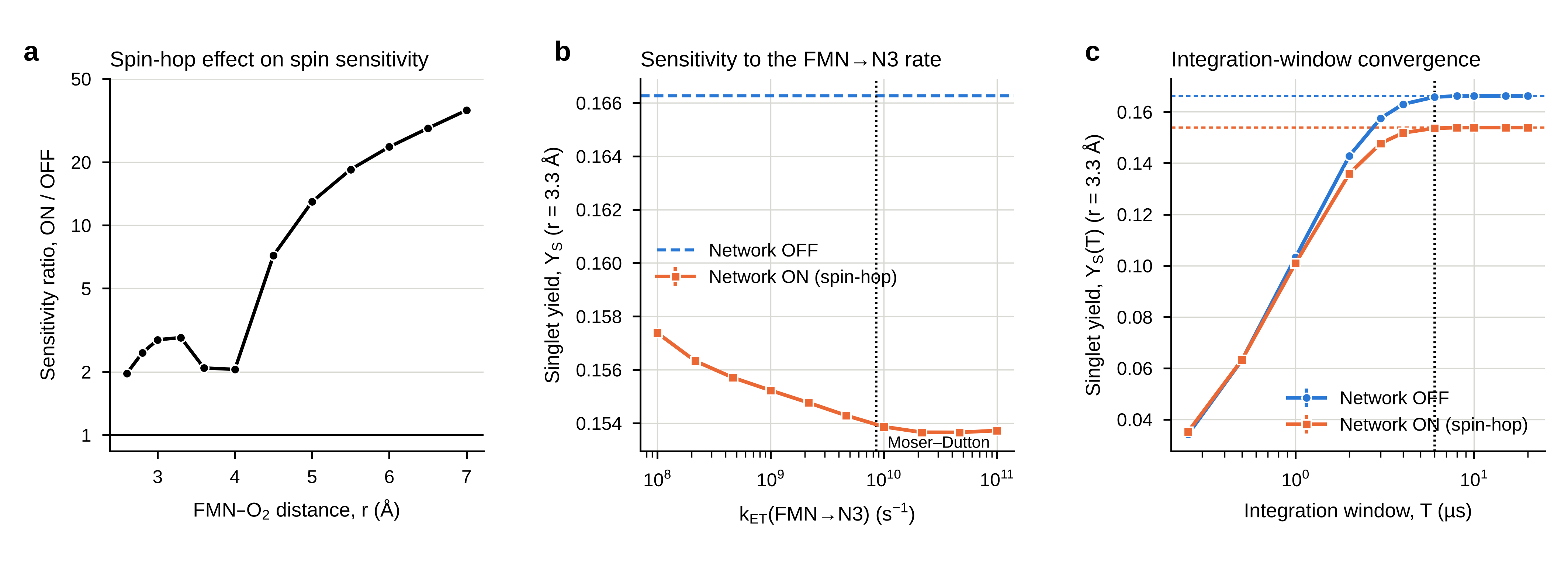

**Supplementary Fig. 4 | Spin-hop model: sensitivity, FMN→N3 rate and integration window.** **a**, Ratio of the combined spin sensitivity, ½(*M*_B_ + *C*_γ_), with network ON versus OFF in the spin-hop model. Both conditions are normalised to the network-OFF maxima so that they can be compared directly. Separations at which the network-OFF magnetic sensitivity is not resolved from zero (8–9 Å) are omitted; 2,000 hyperfine orientations. The charge-resolved model does not reproduce this increase (Supplementary Fig. 3c). **b**, Singlet yield as a function of the FMN→N3 electron-hop rate at *r* = 3.3 Å. Orange, network ON (spin-hop, O₂–FMN–N3); dashed blue line, network OFF. Dotted line, the Moser–Dutton estimate used throughout (8.5 × 10^9^ s^−1^). Across 10^8^–10^11^ s^−1^ the yield varies by 2.4%; 400 orientations. **c**, Singlet yield integrated up to a finite window T, for network OFF (blue) and network ON (orange) at *r* = 3.3 Å. Dashed horizontal lines, the infinite-time yields. Dotted vertical line, the 6 µs window used for time-domain cross-checks, at which both yields lie within 0.3% of their infinite-time values; 200 orientations. *k*_S_ = *k*_T_ = 10^6^ s^−1^. Infinite-time yields were evaluated exactly as *k*_S_ Tr[*P*_S_(−ℒ)^−1^ρ(0)] and finite-window yields as *k*_S_ Tr[*P*_S_ ℒ^−1^(e^(ℒT) − 1)ρ(0)], with the same orientations in every compared condition. Error bars, s.e.m.

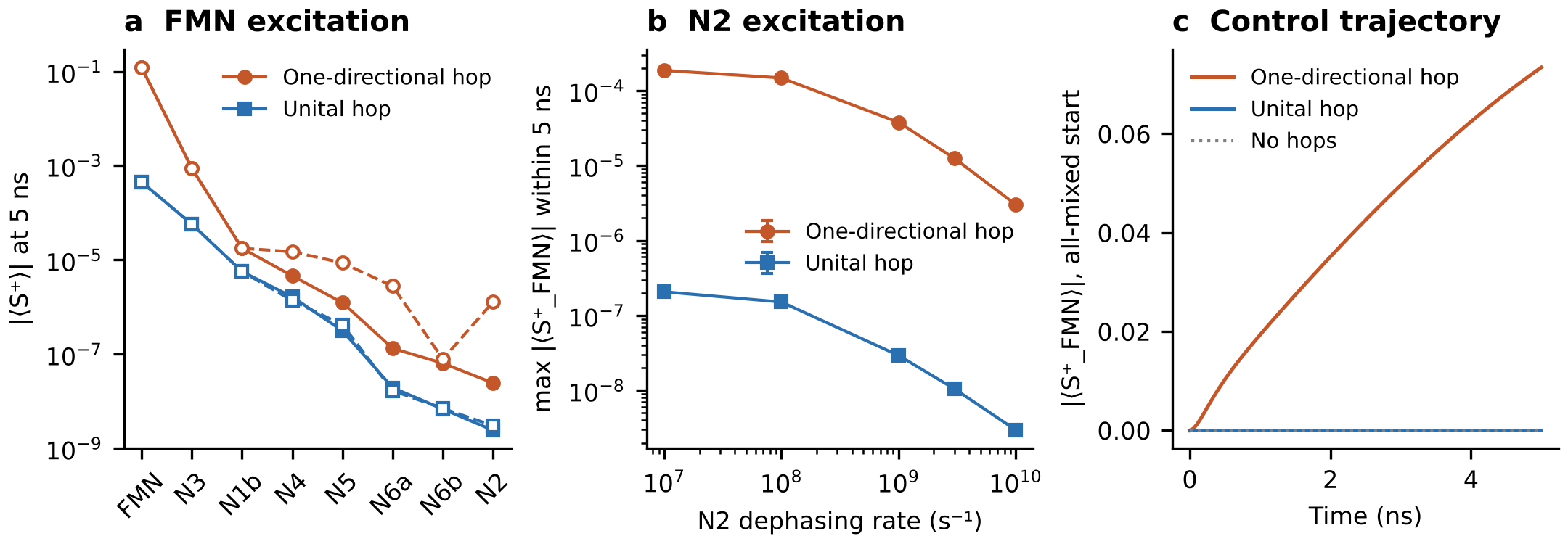

**Supplementary Fig. 5 | One-directional versus unital hop operators in the eight-site coherence-propagation model.** **a**, Baseline-subtracted transverse coherence, |⟨S^+^⟩|, at each centre 5 ns after coherence is injected at FMN, with one-directional flip-flop operators, as used in the radical-pair models of Figs 2d and 3 (orange), and with the unital hops used in Fig. 2a–c, in which each operator √k σ^+^_i_σ^−^_j_ is supplemented by its reverse at the same rate (blue). Filled symbols and solid lines, unknown hop rates (N4→N5, N6a→N6b, N6b→N2) set to zero; open symbols and dashed lines, set to 10^8^ s^−1^. **b**, Maximum coherence reaching FMN within 5 ns after injection at N2, as a function of the N2 dephasing rate. Error bars, s.e.m. **c**, Coherence at FMN in the control trajectory started with every centre maximally mixed (mean of two hyperfine orientations, Supplementary Note 1). The one-directional operator drives the maximally mixed state away from itself and generates coherence at FMN; with unital hops the maximally mixed state is stationary (‖ℒρ_mixed_‖ ≈ 10^−9^ s^−1^) and no coherence appears. The blue data are those shown in Fig. 2a–c. All other parameters as in Fig. 2a–c; 12 random hyperfine orientations, identical in both models.

**Supplementary Table 1 | Geometry, coupling constants and electron-transfer rates of the main-chain links.** Centre-to-centre (*r*_cc_) and edge-to-edge (*r*_edge_) distances, unit vector between centroids (PDB 5XTD frame), point-dipole coupling *d*(*r*_cc_), exchange coupling *J*(*r*_edge_) and forward electron-transfer rate *k*_ET_. FMN→N3, Moser–Dutton relation; other rates from ref. 33; n.a., not available.

| **Link** | ***r*_cc_ (Å)** | ***r*_edge_ (Å)** | **Unit vector** | ***d* (mT)** | ***J* (mT)** | ***k*_ET_ (s^−1^)** |
| --- | --- | --- | --- | --- | --- | --- |
| FMN–N3 | 13.38 | 6.71 | (-0.147, 0.471, -0.870) | 0.775 | 1.1 × 10^−3^ | 8.5 × 10^9^ |
| N3–N1b | 14.57 | 11.42 | (0.443, 0.820, -0.363) | 0.600 | 9.0 × 10^−8^ | 2.9 × 10^6^ |
| N1b–N4 | 13.76 | 11.42 | (-0.913, 0.175, -0.367) | 0.713 | 8.9 × 10^−8^ | 6.4 × 10^4^ |
| N4–N5 | 11.94 | 8.39 | (0.239, -0.182, -0.954) | 1.091 | 3.8 × 10^−5^ | n.a. |
| N5–N6a | 17.20 | 14.82 | (-0.223, 0.188, -0.957) | 0.365 | 1.0 × 10^−10^ | 7.3 × 10^3^ |
| N6a–N6b | 11.45 | 8.36 | (0.649, 0.140, -0.747) | 1.236 | 4.1 × 10^−5^ | n.a. |
| N6b–N2 | 13.73 | 10.33 | (0.583, 0.290, -0.759) | 0.718 | 7.9 × 10^−7^ | n.a. |
| FMN–N1a | 17.40 | 12.24 | (-0.710, -0.646, -0.279) | 0.352 | 1.7 × 10^−8^ | n.a. |

**Supplementary Table 2 | Location of the composite-score maximum under combinations of model choices.** Network-OFF *H*(*r*) on a 0.1 Å grid between 2.5 and 9.0 Å (Supplementary Note 2). Indented rows change one choice relative to the configuration above them.

| **Configuration** | **Peak of *H*(*r*) (Å)** |
| --- | --- |
| All choices as in the earlier analysis | 3.4 |
| … hyperfine field as in this work | 3.4 |
| … O₂–FMN dipolar term included | 3.8 |
| … steric test of this work (FMN excluded) | 4.1 |
| … van der Waals test with FMN atoms also excluded | 5.9 |
| … *M*_B_ as in this work | 2.5 |
| … *C*_γ_ as in this work | 3.3 |
| … normalisation by maximum | 2.5 |
| All choices as in this work | 5.8 |
| … Schulten–Wolynes hyperfine field | 6.3 |
| … no O₂–FMN dipolar term | 5.8 |
| … van der Waals test, FMN atoms not tested | 5.2 |
| … *M*_B_ from 0.05 and 50 mT | 6.2 |
| … *C*_γ_ from 10^6^ and 10^9^ s^−1^ | 9.0 |
| … min–max normalisation | 5.8 |

**Supplementary Table 3 | Key results with the anionic (1.233 mT) and neutral (1.108 mT) flavin hyperfine fields.** Number of paired hyperfine orientations in parentheses. *k*_S_ = *k*_T_ = 10^6^ s^−1^ unless stated (Supplementary Note 3).

| **Quantity** | ***a*_HF_ = 1.233 mT** | ***a*_HF_ = 1.108 mT** |
| --- | --- | --- |
| Δ*Y*_S_ at 3.3 Å, N3 coupling (Fig. 2d; 400) | −0.0123 | −0.0121 |
| Δ*Y*_S_ at 2.6 and 5.0 Å (Fig. 2d; 400) | −0.0042, −0.0155 | −0.0041, −0.0157 |
| Chain-truncation steps at 3.3 Å: +N3, +N1b, +N4 (Fig. 3c; 200) | −0.0124, −0.0458, −0.0053 | −0.0122, −0.0458, −0.0053 |
| Peak of *H*(*r*), FMN excluded: OFF / ON (Fig. 3a; 1,000) | 6.0 / 7.0 Å | 6.5 / 9.0 Å |
| Peak of *H*(*r*), protein only: OFF / ON (Fig. 3a; 1,000) | 5.5 / 6.5 Å | 5.5 / 6.5 Å |
| Threshold midpoints, *k*_S_ = *k*_T_ = 10^6^, 10^7^, 10^8^ s^−1^ (Fig. 3d; 150) | 1.31 × 10^6^, 8.21 × 10^6^, 5.03 × 10^7^ | 1.31 × 10^6^, 8.21 × 10^6^, 5.02 × 10^7^ |
| N1b suppression at the literature rate (Fig. 3d; 150) | 29.7%, 7.0%, 0.69% | 29.8%, 7.0%, 0.69% |
| Sign of the N3 effect on the 5 × 5 rate grid (Supplementary Fig. 1; 400) | 3 negative, 22 positive | 3 negative, 22 positive |
| N3 effect, *k*_S_ = 4 × 10^8^, *k*_T_ = 10^6^ s^−1^, at 3.3 / 6.5 Å (400) | +8.5% / −2.1 ± 0.5% | +9.3% / −2.0 ± 0.6% |

**Supplementary Table 4 | Singlet yields for the model comparison in Fig. 4c with the two exchange conventions.** Orientation-averaged *Y*_S_ (400 paired hyperfine orientations; s.e.m. ≤ 0.006). *Y*_S_^stat^ = *k*_S_/(*k*_S_ + 3*k*_T_). −2*J*, exchange written as −2*J* Ŝ_O₂_·Ŝ_FMN_, as in the charge-resolved model and Fig. 4c; +*J*, exchange written as +*J* Ŝ_O₂_·Ŝ_FMN_, as in the unified spin-hop model of Figs 2d and 3 (Supplementary Note 6).

| ***r* (Å)** | ***k*_S_ (s^−1^)** | ***k*_T_ (s^−1^)** | **Exchange** | ***Y*_S_^stat^** | **Isolated** | **Spin-hop** | **Spin-hop, unital** | **Charge-resolved, N3 held reduced** | **Charge-resolved, N3 re-oxidizable** |
| --- | --- | --- | --- | --- | --- | --- | --- | --- | --- |
| 3.3 | 10^6^ | 10^6^ | −2*J* | 0.2500 | 0.1658 | 0.1533 | 0.2388 | 0.2003 | 0.2297 |
| 3.3 | 10^6^ | 10^6^ | +*J* | 0.2500 | 0.1663 | 0.1540 | 0.2394 | 0.2011 | 0.2298 |
| 3.3 | 10^7^ | 10^7^ | −2*J* | 0.2500 | 0.1432 | 0.1535 | 0.2259 | 0.1499 | 0.1670 |
| 3.3 | 10^7^ | 10^7^ | +*J* | 0.2500 | 0.1432 | 0.1536 | 0.2263 | 0.1501 | 0.1672 |
| 3.3 | 10^8^ | 10^8^ | −2*J* | 0.2500 | 0.0627 | 0.1137 | 0.2184 | 0.0644 | 0.0684 |
| 3.3 | 10^8^ | 10^8^ | +*J* | 0.2500 | 0.0627 | 0.1137 | 0.2185 | 0.0644 | 0.0685 |
| 3.3 | 4 × 10^8^ | 10^6^ | −2*J* | 0.9926 | 0.3680 | 0.4021 | 0.8391 | 0.5207 | 0.8310 |
| 3.3 | 4 × 10^8^ | 10^6^ | +*J* | 0.9926 | 0.3785 | 0.4123 | 0.8462 | 0.5298 | 0.8320 |
| 3.3 | 10^6^ | 10^8^ | −2*J* | 0.0033 | 0.0032 | 0.0057 | 0.0033 | 0.0032 | 0.0029 |
| 3.3 | 10^6^ | 10^8^ | +*J* | 0.0033 | 0.0032 | 0.0057 | 0.0033 | 0.0032 | 0.0029 |
| 6.5 | 10^6^ | 10^6^ | −2*J* | 0.2500 | 0.2120 | 0.2022 | 0.2442 | 0.2257 | 0.2358 |
| 6.5 | 10^6^ | 10^6^ | +*J* | 0.2500 | 0.2121 | 0.2022 | 0.2442 | 0.2257 | 0.2358 |
| 6.5 | 10^7^ | 10^7^ | −2*J* | 0.2500 | 0.1596 | 0.1691 | 0.2305 | 0.1670 | 0.1792 |
| 6.5 | 10^7^ | 10^7^ | +*J* | 0.2500 | 0.1596 | 0.1691 | 0.2305 | 0.1671 | 0.1793 |
| 6.5 | 10^8^ | 10^8^ | −2*J* | 0.2500 | 0.0745 | 0.1331 | 0.2152 | 0.0767 | 0.0804 |
| 6.5 | 10^8^ | 10^8^ | +*J* | 0.2500 | 0.0745 | 0.1331 | 0.2152 | 0.0767 | 0.0804 |
| 6.5 | 4 × 10^8^ | 10^6^ | −2*J* | 0.9926 | 0.8245 | 0.8215 | 0.9070 | 0.8495 | 0.9048 |
| 6.5 | 4 × 10^8^ | 10^6^ | +*J* | 0.9926 | 0.8248 | 0.8217 | 0.9070 | 0.8497 | 0.9049 |
| 6.5 | 10^6^ | 10^8^ | −2*J* | 0.0033 | 0.0032 | 0.0053 | 0.0033 | 0.0032 | 0.0030 |
| 6.5 | 10^6^ | 10^8^ | +*J* | 0.0033 | 0.0032 | 0.0053 | 0.0033 | 0.0032 | 0.0030 |

**Supplementary Table 5 | Charge-resolved singlet yields as a function of the FMN→N3 driving force (*k*_T_ = 10^6^ s^−1^).** *k*_f_ from the Moser–Dutton relation and *k*_b_ from detailed balance at 310 K; 150 paired hyperfine orientations (Supplementary Note 6). Isolated-pair yields for comparison (ΔG-independent): 0.166 and 0.371 at 3.3 Å, 0.214 and 0.833 at 6.5 Å, for *k*_S_ = 10^6^ and 4 × 10^8^ s^−1^, respectively.

| **ΔG (eV)** | ***k*_f_ (s^−1^)** | ***k*_b_ (s^−1^)** | ***r* (Å)** | ***k*_S_ (s^−1^)** | **Reduced chain, N3 held reduced** | **Reduced chain, N3 re-oxidizable** | **Oxidized N3, onward 6.4 × 10^4^ s^−1^** |
| --- | --- | --- | --- | --- | --- | --- | --- |
| −0.05 | 5.23 × 10^9^ | 8.05 × 10^8^ | 3.3 | 10^6^ | 0.2004 | 0.2297 | 0.1765 |
| −0.05 | 5.23 × 10^9^ | 8.05 × 10^8^ | 3.3 | 4 × 10^8^ | 0.5223 | 0.8310 | 0.9873 |
| −0.05 | 5.23 × 10^9^ | 8.05 × 10^8^ | 6.5 | 10^6^ | 0.2263 | 0.2360 | 0.1765 |
| −0.05 | 5.23 × 10^9^ | 8.05 × 10^8^ | 6.5 | 4 × 10^8^ | 0.8562 | 0.9074 | 0.9873 |
| −0.09 | 8.55 × 10^9^ | 2.94 × 10^8^ | 3.3 | 10^6^ | 0.2004 | 0.2297 | 0.0874 |
| −0.09 | 8.55 × 10^9^ | 2.94 × 10^8^ | 3.3 | 4 × 10^8^ | 0.5223 | 0.8312 | 0.9733 |
| −0.09 | 8.55 × 10^9^ | 2.94 × 10^8^ | 6.5 | 10^6^ | 0.2263 | 0.2360 | 0.0874 |
| −0.09 | 8.55 × 10^9^ | 2.94 × 10^8^ | 6.5 | 4 × 10^8^ | 0.8562 | 0.9075 | 0.9733 |
| −0.164 | 1.88 × 10^10^ | 4.05 × 10^7^ | 3.3 | 10^6^ | 0.2004 | 0.2297 | 0.0081 |
| −0.164 | 1.88 × 10^10^ | 4.05 × 10^7^ | 3.3 | 4 × 10^8^ | 0.5223 | 0.8313 | 0.7629 |
| −0.164 | 1.88 × 10^10^ | 4.05 × 10^7^ | 6.5 | 10^6^ | 0.2263 | 0.2360 | 0.0081 |
| −0.164 | 1.88 × 10^10^ | 4.05 × 10^7^ | 6.5 | 4 × 10^8^ | 0.8562 | 0.9075 | 0.7629 |
| −0.2 | 2.61 × 10^10^ | 1.46 × 10^7^ | 3.3 | 10^6^ | 0.2004 | 0.2297 | 0.0022 |
| −0.2 | 2.61 × 10^10^ | 1.46 × 10^7^ | 3.3 | 4 × 10^8^ | 0.5223 | 0.8313 | 0.4614 |
| −0.2 | 2.61 × 10^10^ | 1.46 × 10^7^ | 6.5 | 10^6^ | 0.2263 | 0.2360 | 0.0022 |
| −0.2 | 2.61 × 10^10^ | 1.46 × 10^7^ | 6.5 | 4 × 10^8^ | 0.8562 | 0.9076 | 0.4614 |

**Supplementary Table 6 | Recombination-rate grid of the charge-resolved network effect at *r* = 3.3 Å (Fig. 4d).** Orientation-averaged *Y*_S_; 150 paired hyperfine orientations per point. Change, percentage change in *Y*_S_ produced by the charge-resolved network with N3 re-oxidizable, relative to the isolated pair (as plotted in Fig. 4d). Rule, whether this change has the sign of *Y*_S_^stat^ − *Y*_S_^iso^. Spin-hop, one-directional spin-hop model for comparison.

| ***k*_S_ (s^−1^)** | ***k*_T_ (s^−1^)** | ***Y*_S_^stat^** | **Isolated** | **Charge-resolved, N3 held reduced** | **Charge-resolved, N3 re-oxidizable** | **Change (%)** | **Rule** | **Spin-hop** |
| --- | --- | --- | --- | --- | --- | --- | --- | --- |
| 10^5^ | 10^5^ | 0.2500 | 0.1819 | 0.2401 | 0.2475 | +36.1 | yes | 0.1007 |
| 10^5^ | 10^6^ | 0.0323 | 0.0303 | 0.0313 | 0.0300 | −0.9 | no | 0.0250 |
| 10^5^ | 10^7^ | 0.0033 | 0.0033 | 0.0033 | 0.0029 | −12.9 | no | 0.0035 |
| 10^5^ | 10^8^ | 0.0003 | 0.0003 | 0.0003 | 0.0003 | −9.5 | no | 0.0006 |
| 10^5^ | 10^9^ | 0.0000 | 0.0000 | 0.0000 | 0.0000 | −8.5 | no | 0.0003 |
| 10^6^ | 10^5^ | 0.7692 | 0.3587 | 0.6826 | 0.7606 | +112.1 | yes | 0.3647 |
| 10^6^ | 10^6^ | 0.2500 | 0.1660 | 0.2004 | 0.2297 | +38.4 | yes | 0.1534 |
| 10^6^ | 10^7^ | 0.0323 | 0.0294 | 0.0297 | 0.0269 | −8.6 | no | 0.0310 |
| 10^6^ | 10^8^ | 0.0033 | 0.0032 | 0.0032 | 0.0029 | −9.0 | no | 0.0057 |
| 10^6^ | 10^9^ | 0.0003 | 0.0003 | 0.0003 | 0.0003 | −8.2 | no | 0.0026 |
| 10^7^ | 10^5^ | 0.9709 | 0.4050 | 0.8372 | 0.9548 | +135.7 | yes | 0.5004 |
| 10^7^ | 10^6^ | 0.7692 | 0.3015 | 0.4369 | 0.6720 | +122.9 | yes | 0.3171 |
| 10^7^ | 10^7^ | 0.2500 | 0.1432 | 0.1499 | 0.1670 | +16.7 | yes | 0.1536 |
| 10^7^ | 10^8^ | 0.0323 | 0.0233 | 0.0235 | 0.0222 | −4.7 | no | 0.0419 |
| 10^7^ | 10^9^ | 0.0033 | 0.0025 | 0.0025 | 0.0023 | −5.7 | no | 0.0204 |
| 10^8^ | 10^5^ | 0.9970 | 0.4769 | 0.8607 | 0.9769 | +104.8 | yes | 0.5742 |
| 10^8^ | 10^6^ | 0.9709 | 0.3379 | 0.5004 | 0.8158 | +141.4 | yes | 0.3660 |
| 10^8^ | 10^7^ | 0.7692 | 0.2343 | 0.2525 | 0.3608 | +54.0 | yes | 0.2554 |
| 10^8^ | 10^8^ | 0.2500 | 0.0627 | 0.0644 | 0.0685 | +9.2 | yes | 0.1137 |
| 10^8^ | 10^9^ | 0.0323 | 0.0076 | 0.0078 | 0.0077 | +1.2 | yes | 0.0628 |
| 10^9^ | 10^5^ | 0.9997 | 0.7351 | 0.8923 | 0.9798 | +33.3 | yes | 0.7838 |
| 10^9^ | 10^6^ | 0.9970 | 0.4227 | 0.5519 | 0.8372 | +98.1 | yes | 0.4587 |
| 10^9^ | 10^7^ | 0.9709 | 0.2601 | 0.2799 | 0.4174 | +60.5 | yes | 0.2886 |
| 10^9^ | 10^8^ | 0.7692 | 0.0765 | 0.0789 | 0.0964 | +26.0 | yes | 0.1420 |
| 10^9^ | 10^9^ | 0.2500 | 0.0097 | 0.0100 | 0.0105 | +9.0 | yes | 0.0824 |
