## Supplementary figures and images for "Kinetic gating by an iron–sulfur chain modulates flavin radical-pair spin chemistry in mitochondrial Complex I"

### supplefigure1

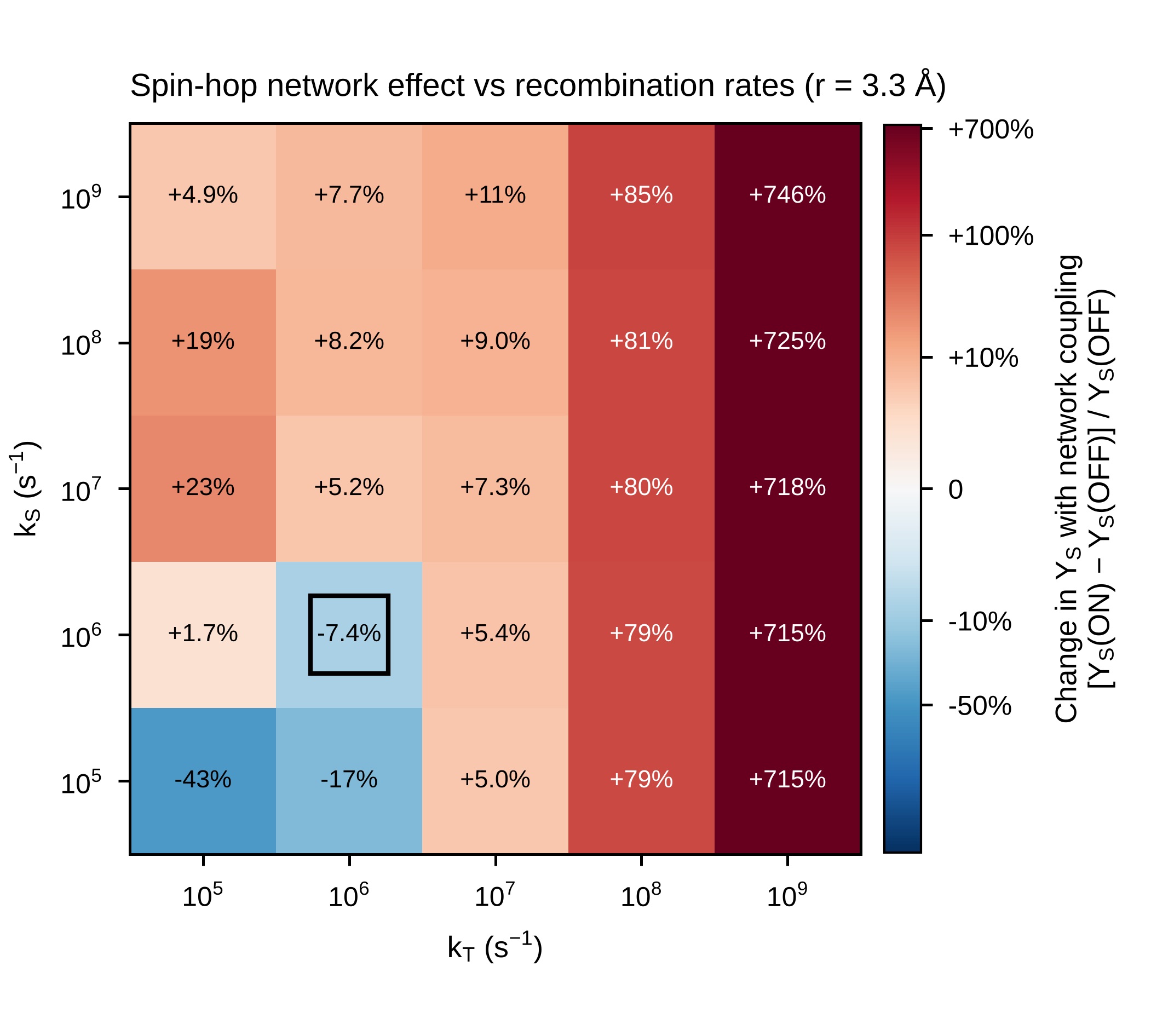

### supplefigure2

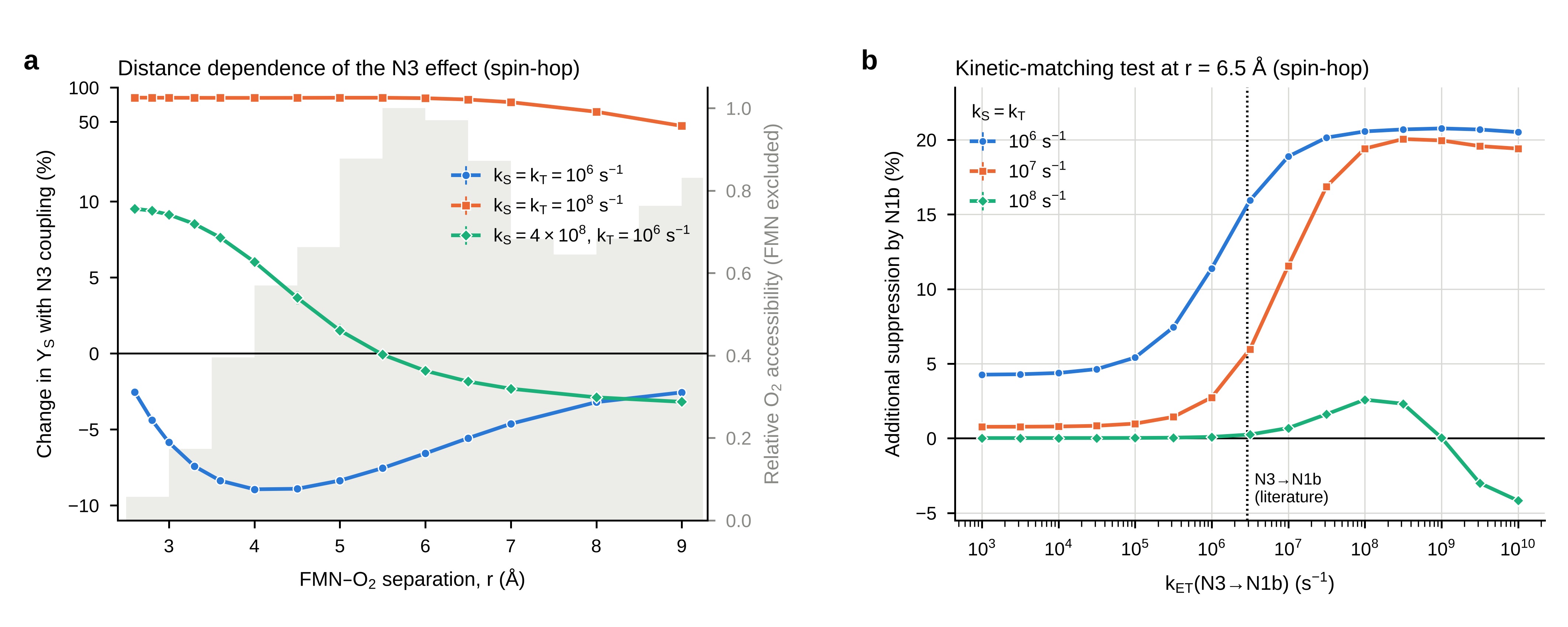

### supplefigure3

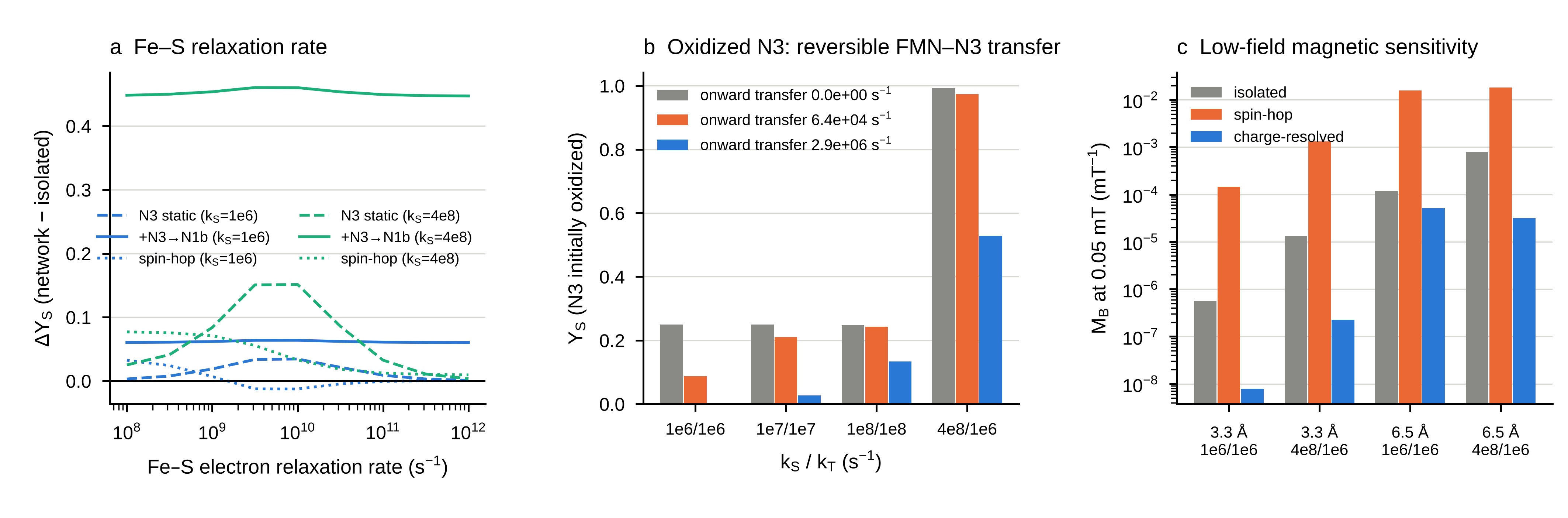

### supplefigure4

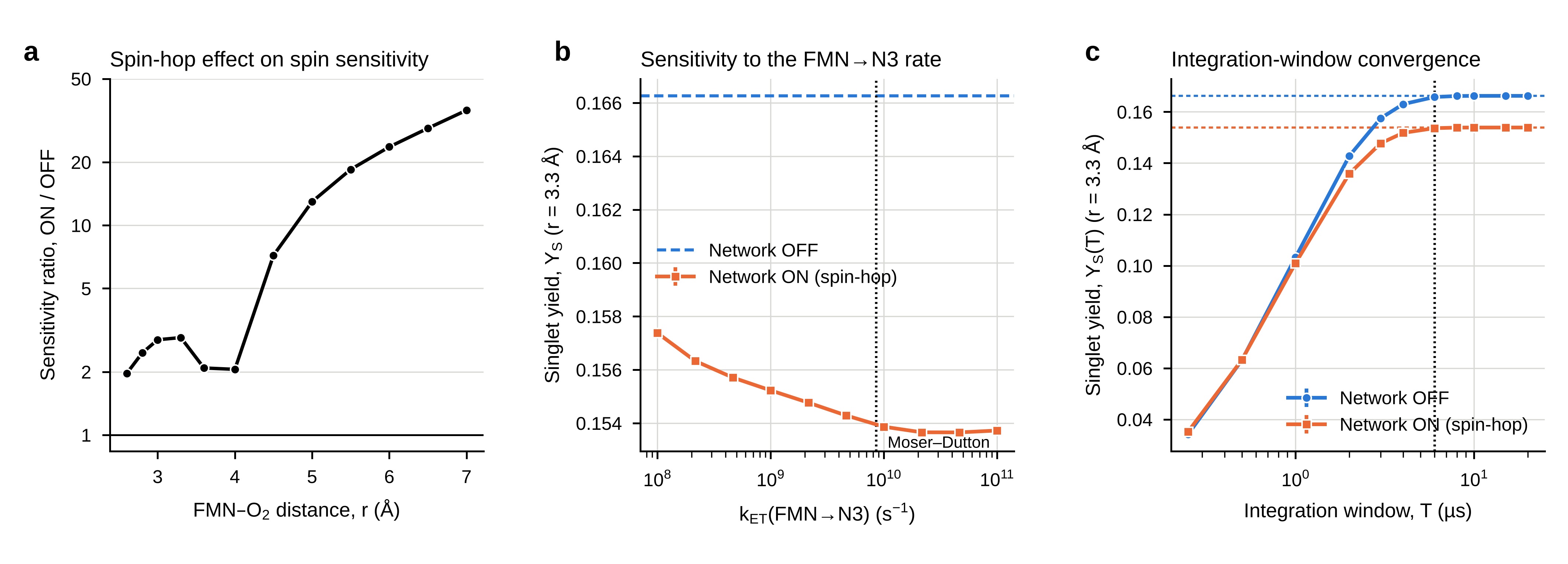

### supplefigure5

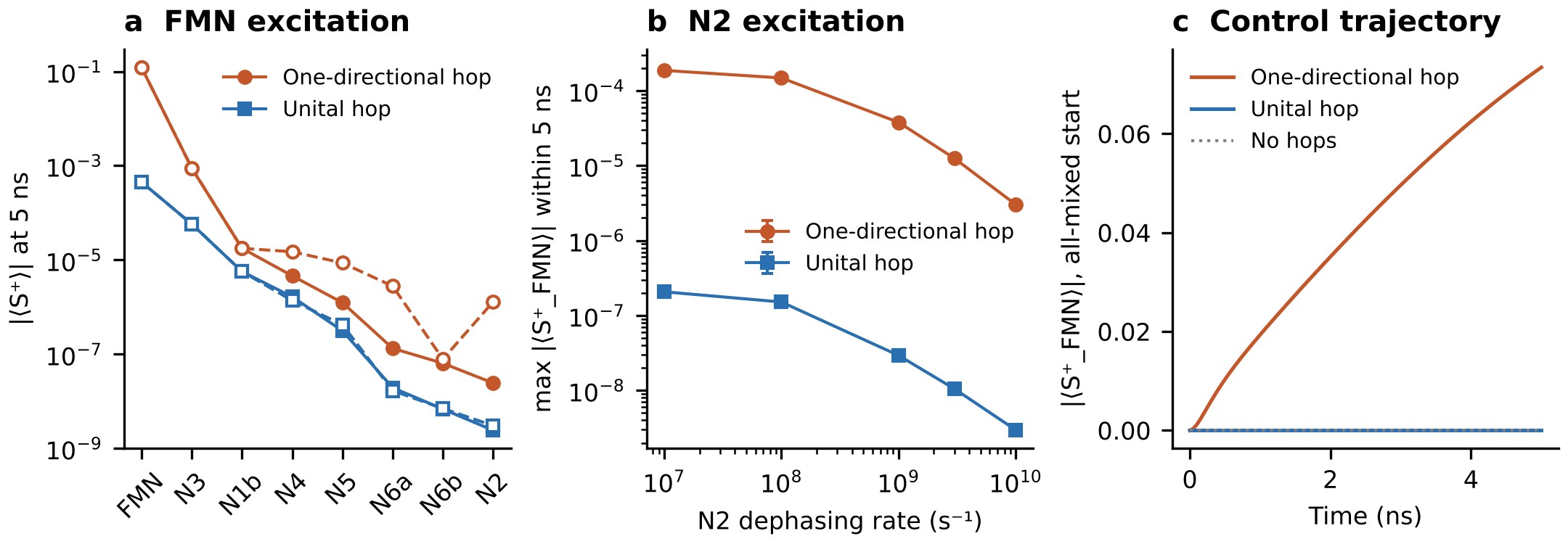
